# Eco1-dependent cohesin acetylation anchors chromatin loops and cohesion to define functional meiotic chromosome domains

**DOI:** 10.1101/2021.09.24.461725

**Authors:** Rachael E. Barton, Lucia F. Massari, Daniel Robertson, Adele L. Marston

## Abstract

Cohesin organizes the genome by forming intra-chromosomal loops and inter-sister chromatid linkages. During gamete formation by meiosis, chromosomes are reshaped to support crossover recombination and two consecutive rounds of chromosome segregation. Here we show that Eco1 acetyltransferase positions both chromatin loops and sister chromatid cohesion to organize meiotic chromosomes into functional domains in budding yeast. Eco1 acetylates the Smc3 cohesin subunit in meiotic S phase to establish chromatin boundaries, independently of DNA replication. Boundary formation by Eco1 is critical for prophase exit and for the maintenance of cohesion until meiosis II, but is independent of the ability of Eco1 to antagonize the cohesin-release factor, Wpl1. Conversely, prevention of cohesin release by Wpl1 is essential for centromeric cohesion, kinetochore monoorientation and co-segregation of sister chromatids in meiosis I. Our findings establish Eco1 as a key determinant of chromatin boundaries and cohesion positioning, revealing how local chromosome structuring directs genome transmission into gametes.

## Introduction

The cohesin complex defines genome architecture to support DNA repair, gene expression and chromosome segregation (Davidson and Peters, 2021). Core cohesin is a DNA translocase comprising a V-shaped heterodimer of two structural maintenance of chromosomes proteins, Smc1 and Smc3, whose two ATPase heads are connected by a kleisin subunit. Cohesin folds the genome through ATP-dependent extrusion of intra-molecular DNA loops. Cohesin also entraps newly replicated sister chromatids within its tripartite structure, to establish the cohesion needed for chromosome segregation. Loop extrusion and cohesion are biochemically distinct and dependent on cohesin accessory proteins, although the mechanisms are not completely understood (Srinivasan et al., 2018). Loading of cohesin onto DNA requires the Scc2-Scc4 (NIPBL-MAU2 in mammals) complex, which also drives loop extrusion (Ciosk et al., 2000; Davidson et al., 2019; Petela et al., 2018; Srinivasan et al., 2019). Chromosomal cohesin is destabilized by the cohesin release factor, Wpl1/Rad61 (WAPL in mammals), an activity that is counteracted by acetylation of cohesin’s Smc3 subunit by the Eco1 acetyltransferase (Ben-Shahar et al., 2008; Unal et al., 2008).

Cohesin-mediated loops are positioned by boundary elements. In mammalian interphase cells, the insulation protein CTCF anchors cohesin at the sites of long-range interactions (Haarhuis et al., 2017; Rao et al., 2017; Schwarzer et al., 2017; Wutz et al., 2017). In yeast, CTCF is absent but chromosomes are nevertheless organized into cohesin-dependent loops in mitosis and convergent genes, which are known to accumulate cohesin, are found at loop boundaries (Costantino et al., 2020; Lazar-Stefanita et al., 2017; Lengronne et al., 2004; Paldi et al., 2020; Schalbetter et al., 2017). Budding yeast pericentromeres provide an exemplary model of how a functional chromosome domain is folded. Cohesin loaded at centromeres extrudes loops on both sides until it is stalled by convergent genes at flanking pericentromere boundaries, thereby establishing a structure that facilitates chromosome segregation in mitosis (Paldi et al., 2020). Loop size and position are also controlled by cohesin regulators. Wpl1 restricts loop size, but does not affect their positioning (Costantino et al., 2020; Dauban et al., 2020; Haarhuis et al., 2017). Eco1 also limits long-range interactions, but additionally affects loop positioning (Dauban et al., 2020). Whether this is a direct effect, through Eco1-mediated acetylation and inhibition of loop-extruding cohesin, or indirect, as a result of acetylated cohesive cohesin forming a barrier to translocating cohesin, is unclear. Cells lacking Eco1 are inviable due to cohesion defects, which can be rescued by removal of Wpl1 to restore viability (Ben-Shahar et al., 2008; Rowland et al., 2009; Sutani et al., 2009; Unal et al., 2008).

However, cells lacking both Wpl1 and Eco1 show an additive increase in long-range interactions (Dauban et al., 2020). Together, these observations suggested that loop positioning is not essential for viability. Consistent with its essential function in establishing cohesion, in mitotically growing cells acetylation of Smc3 by Eco1 is coupled with DNA replication (Beckouët et al., 2010; Ben-Shahar et al., 2008). In mammals, ESCO2 similarly acetylates SMC3 during S phase to establish cohesion (Alomer et al., 2017) while, during interphase, an additional family member, ESCO1, is active and contributes to boundary formation in chromatin looping (Wutz et al., 2020). Together, these observations indicate that Eco1-dependent cohesin acetylation can both position loops and maintain cohesion.

During gamete formation by meiosis, chromosomes undergo extensive restructuring, underpinned by cohesin-dependent chromosome looping and cohesion. In many organisms a meiosis-specific kleisin, Rec8, enables functions that cannot be carried out by canonical kleisin, Scc1/Rad21 (Severson et al., 2009; Tachibana-Konwalski et al., 2010; Toth et al., 2000; Yokobayashi et al., 2003). During meiotic prophase, chromosomes comprise a dense array of chromatin loops emanating from cohesin-rich axes that are zipped together in homologous pairs by a central core - the synaptonemal complex (Cahoon and Hawley, 2016). Budding yeast Rec8-cohesin anchors loops at their base (Muller et al., 2018; Schalbetter et al., 2019) and supports crossover recombination and DNA repair to allow prophase exit (Klein et al., 1999). Between prophase I and metaphase I, Wpl1 removes a fraction of chromosomal Rec8 (Challa et al., 2016, 2019). WAPL also promotes meiotic release of cohesin in *Arabidopsis thaliana* and *Caenorhabditis elegans*, and reduces loop number in mouse oocytes, although the kleisin target differs between organisms. (Crawley et al., 2016; De et al., 2014; Silva et al., 2020). Following prophase exit, two distinct meiotic divisions ensue (reviewed in (Duro and Marston, 2015)). During meiosis I, sister kinetochores are monooriented to ensure sister chromatid co-segregation. In budding yeast, the monopolin complex fuses sister kinetochores, while in fission yeast and mammals centromeric Rec8-cohesin directs sister chromatid co-segregation (Chelysheva et al., 2005; Monje-Casas et al., 2007; Parra et al., 2004; Sakuno et al., 2009; Severson et al., 2009). Homolog segregation at meiosis I is triggered by separase-dependent cleavage of Rec8 on chromosome arms, while pericentromeric cohesin is retained and cleaved only at meiosis II to allow sister chromatid segregation. How cohesin-dependent loop formation and cohesion are spatially and temporally regulated to establish distinct functional chromosome domains in meiosis remains unclear.

Here we identify Eco1 acetyltransferase as a key determinant of localized meiotic chromosome structure in budding yeast. In meiosis, Eco1 acetylates Smc3 independently of DNA replication and is critical for viability, even in the absence of Wpl1. Eco1 counteracts Wpl1 to allow centromeric cohesion establishment and thereby kinetochore monoorientation. In contrast, arm cohesion and prophase exit all require an Eco1 function other than Wpl1 antagonism. While Eco1 and Wpl1 independently restrict chromatin loop size in prophase I, only Eco1 is critical for boundary formation, notably at pericentromere borders. We propose that cohesin acetylation by Eco1 traps both loop extruding and cohesive cohesin complexes at boundaries to define a chromosome architecture that is essential for meiotic recombination and chromosome segregation.

## Results

### Eco1 acetylates cohesin during meiotic S phase, independently of DNA replication

To determine the timing of Eco1-dependent Smc3 acetylation during meiosis, wild type cells carrying functional *ECO1-6HIS-3FLAG* (Figure S1A), were released from a pre-meiotic S phase block (Berchowitz et al., 2013) allowing synchronous DNA replication and nuclear divisions (Figure 1A and B). Eco1 levels increased after meiotic entry, were maximal during meiotic S phase and declined around the time of prophase exit, while Smc3-K112,K113 acetylation (henceforth Smc3-Ac), detected using a verified antibody (Figure S1B and C), accumulated after Eco1 appearance, persisted throughout the meiotic divisions and declined during meiosis II (Figure 1C).

**Figure 1.**
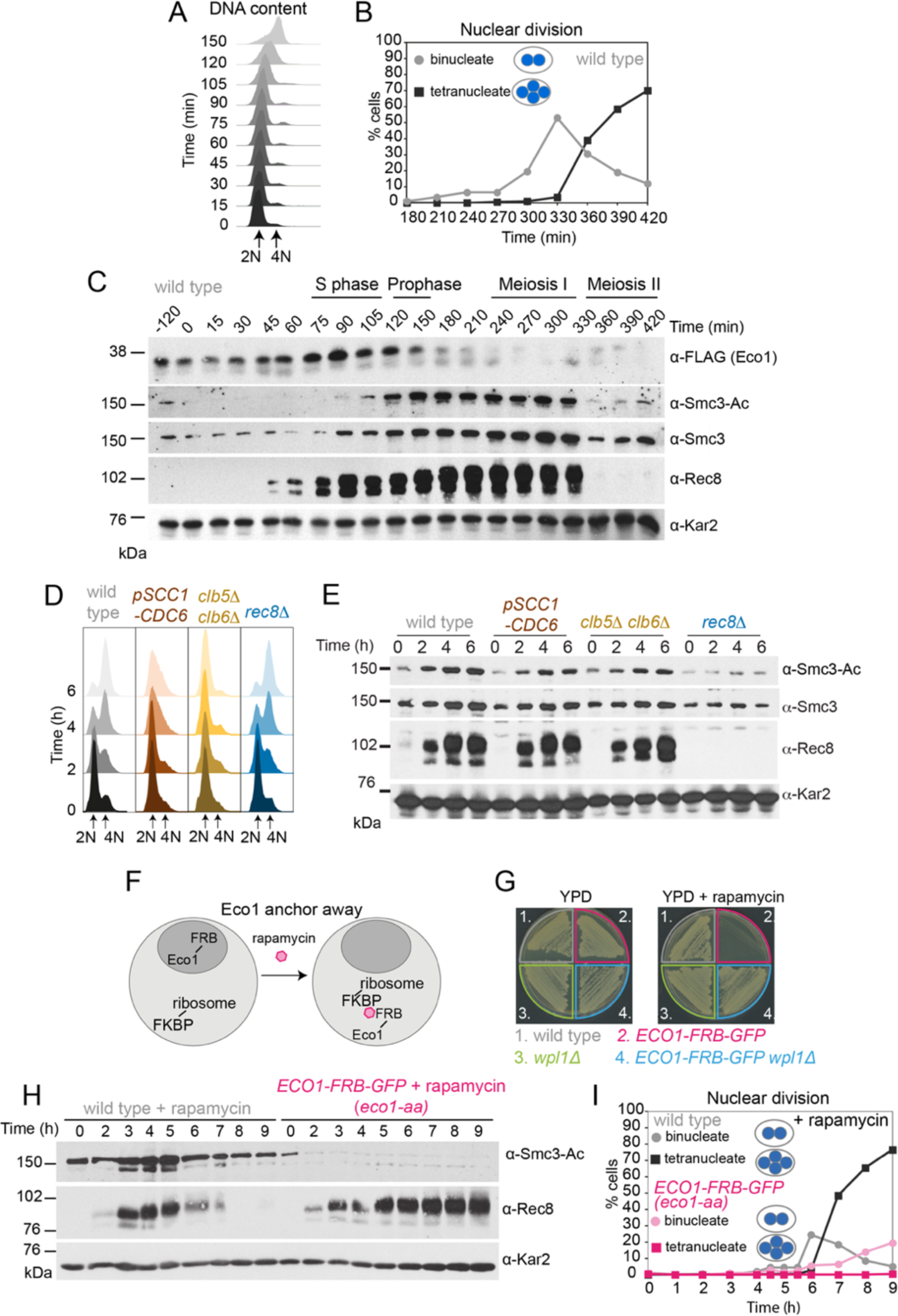
Eco1-dependent acetylation of Smc3-K112,113 occurs in meiotic S phase, independently of DNA replication. (A-C) Smc3-Ac is deposited in S phase, following Eco1 production. Wild type (strain AM21574) carrying *ECO1-6HIS-3FLAG* and *pCUP1-IME1 pCUP1-IME4* was released from a pre-meiotic S phase block 120 min after sporulation induction by addition of 25 μM CuSO_4_. (A) S phase completion (4N) was monitored by flow cytometry. (B) The percentages of bi- and tetra-nucleate cells were scored at the indicated time points to monitor meiosis I and II nuclear division, respectively (n=200). (C) Western immunoblot shows the total cellular levels of Eco1-6HIS-3FLAG (α-FLAG), Smc3-Ac (α-Smc3-K112,113-Ac), Smc3 (α-Smc3) and Rec8 (α-Rec8) with Kar2 as a loading control (α-Kar2). (D and E) Bulk DNA replication is not essential for Smc3-Ac. Wild type (AM11633), *pSCC1-CDC6* (AM28842), *clb5Δ clb6Δ* (AM28841) and *rec8Δ* (AM28843) cells carrying *ndt80Δ* were induced to sporulate and allowed to arrest in prophase I. (D) Flow cytometry shows DNA content. (E) Western immunoblot shows total cellular levels of Smc3-Ac (α-Smc3-K112,113-Ac), Smc3 (*α*-Smc3), Rec8 (*α*-Rec8) and Kar2 loading control (*α*-Kar2). (F) Schematic of the anchor-away system used to deplete Eco1 from the nucleus (*eco1-aa*). (G) The lethality of *ECO1* anchor-away is rescued by deletion of *WPL1*. Haploid wild type (AM13762), *ECO1-FRB-GFP* (AM22004), *wpl1Δ* (AM22440), and *ECO1-FRB-GFP wpl1Δ* (AM22981) strains of the anchor-away background (*RPL13A-FKBP12, fpr1Δ, tor1-1*) were plated on YPD or YPD+1 μM rapamycin. (H and I) Eco1 is essential for meiotic progression. Anchoring-away Eco1-FRB-GFP reduces acetylation of Smc3-K112,K113, impairs cleavage of Rec8 and reduces nuclear divisions. Anchor-away wild type (AM25532) and *ECO1-FRB-GFP* (AM22034) cells were induced to sporulate in the presence of 1 μM rapamycin. (H) Western immunoblot of whole cell extracts showing Smc3-Ac (α-Smc3-K112,K113-Ac), Rec8 (α-Rec8), and Kar2 loading control (α-Kar2). (I) The percentages of bi- and tetra-nucleate cells were scored after DAPI-staining at the indicated timepoints (n = 200 cells/time point).

To test whether DNA replication is required for Smc3 acetylation in meiosis, we analyzed *pSCC1-CDC6* and *clb5Δ clb6Δ* cells that fail to assemble or fire, pre - replicative complexes, respectively, resulting in little or no replication in pre-meiotic S phase (Figure 1D; (Brar et al., 2009; Stuart and Wittenberg, 1998)). Surprisingly, Smc3-Ac appeared in *pSCC1-CDC6* and *clb5Δ clb6Δ* meiotic cells with comparable timing to wild type and was only modestly reduced (Figure 1E). In contrast, Smc3-Ac was greatly diminished in cells lacking the meiotic cohesin kleisin subunit (*rec8Δ*) (Figure 1E). Because DNA double-strand breaks trigger Eco1-dependent cohesion establishment in mitotic cells (Strom et al., 2007; Unal et al., 2007), we tested whether Spo11 endonuclease-induced meiotic double strand breaks were required for Smc3-Ac. However, preventing double strand break formation (*spo11Δ*) did not reduce Smc3-Ac levels whether chromosomes were replicated or not (Figure S1D and E). We conclude that cohesin acetylation on its Smc3 subunit during meiotic S phase occurs independently of DNA replication and programmed double strand break formation.

### Eco1 is essential for meiosis, even in the absence of WPL1

Eco1 is essential for vegetative growth and *eco1Δ* viability is restored by deletion of *WPL1* (*eco1Δ wpl1Δ* cells) (Ben-Shahar et al., 2008). To examine Eco1 function specifically in meiosis we fused it to the FKBP-rapamycin binding domain (FRB) to allow rapamycin-induced anchoring out of the nucleus (Haruki et al., 2008) (Figure 1F; see also materials and methods). Growth of *ECO1-FRB-GFP* cells was inhibited in the presence of rapamycin, but restored by deletion of *WPL1* (Figure 1G), consistent with successful anchor-away (Ben-Shahar et al., 2008; Sutani et al., 2009; Unal et al., 2008). Upon induction of meiosis in the presence of rapamycin, Smc3-Ac was undetectable in *ECO1-FRB-GFP* cells and full-length Rec8 persisted long after its expected time of degradation (Figure 1H). Furthermore, only a minor fraction of *ECO1-FRB-GFP* cells completed the first meiotic division (Figure 1I). Even in the absence of rapamycin, *ECO1-FRB-GFP* cells showed defects in meiotic progression, Rec8 degradation and Smc3-Ac (Figure S1F and G), despite supporting vegetative growth (Figure 1G). Therefore, the FRB-GFP tag on Eco1 specifically affects meiosis. Henceforth, *ECO1-FRB-GFP* cells were induced to sporulate in the presence of rapamycin and are denoted *eco1-aa.* We conclude that Eco1 acetylates Smc3 on residues K112,113 during S phase of meiosis, and that Eco1 is required for efficient meiotic division.

We reasoned that Eco1 activity early in meiosis may be required to counter Wpl1-dependent cohesin destabilization during later stages of meiosis. If this is the case, deletion of Wpl1 may overcome the meiotic arrest of *eco1-aa* cells. Wpl1 promotes the non-proteolytic removal of cohesin between meiotic prophase and metaphase I (Challa et al., 2016, 2019). Prior to meiotic S phase, functional Wpl1-6HA gains an activating phosphorylation (Challa et al., 2019) and its overall levels increase (Figure 2A-C; Figure S2A-C). Subsequently, and reminiscent of the sequential loss of cohesin from chromosome arms and pericentromeres, Wpl1 undergoes step-wise degradation during meiosis I and II (Figure S2B and C). To test the idea that Eco1 allows meiotic progression by countering Wpl1, we sought to measure spore formation and viability (Figure 2D; Figure S2D-F). However, deletion of *WPL1* in the *eco1-aa* background only slightly increased the formation, but not the viability, of spores, while *wpl1Δ* cells showed a small decrease in spore viability, as reported ((Challa et al., 2016); Figure 2D; Figure S2D,). Consistently, *WPL1* deletion did not restore nuclei division to *eco1-aa* cells (Figure S2E). Sporulating *ECO1-FRB-GFP* cells in the absence of rapamycin increased sporulation efficiency, but only slightly improved viability (Figure S2D-F). Therefore, in contrast to vegetative cells, Eco1 is essential for meiosis, even in the absence of Wpl1.

**Figure 2.**
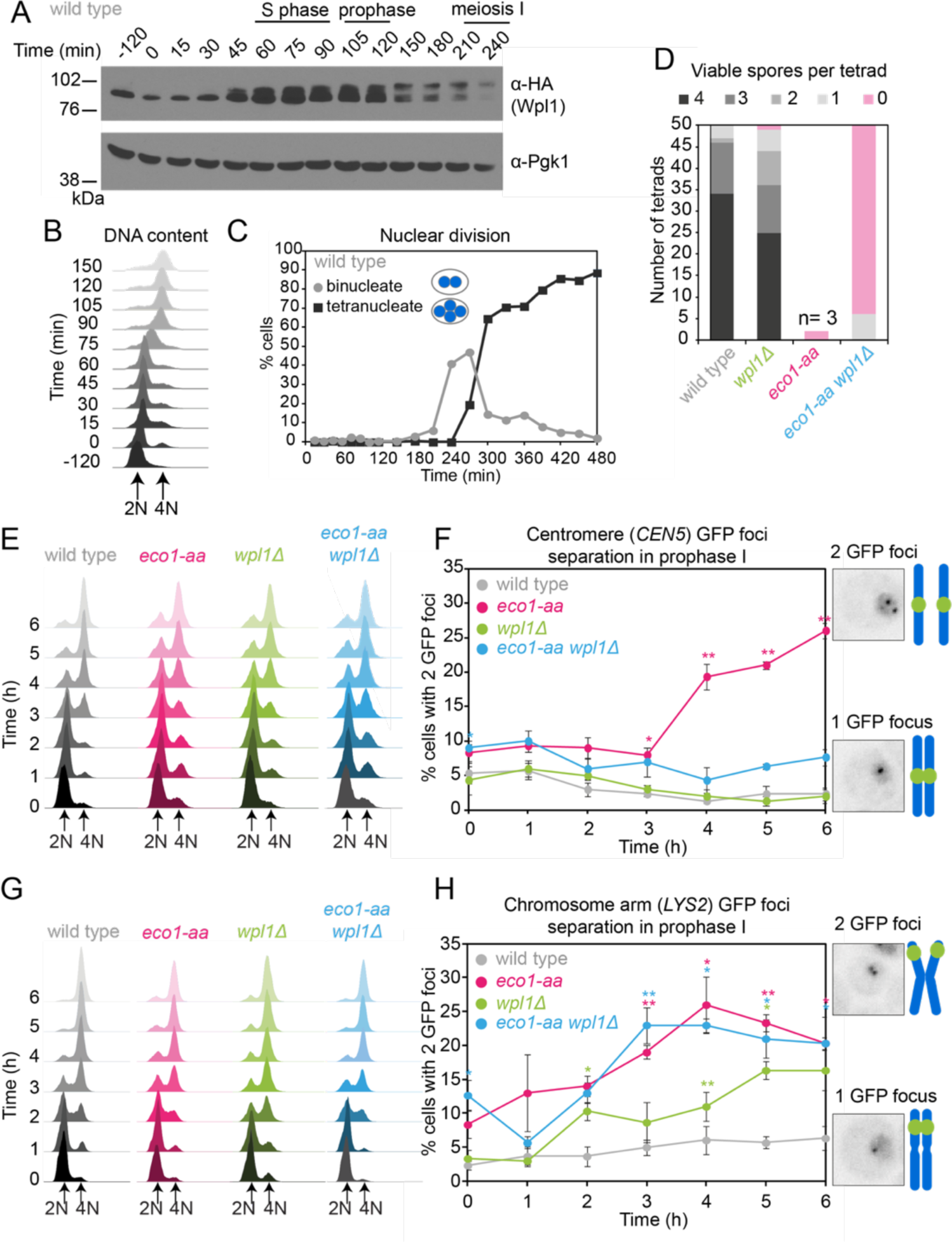
Counteracting Wpl1 is not the only essential role of Eco1 in meiosis. (A – C) Wpl1 is most abundant during meiotic S phase and prophase. Wild type (AM20916) carrying *WPL1-6HA* and *pCUP1-IME1 pCUP1-IME4* was induced to undergo synchronous meiotic S phase as described in Figure 1A. (A) Western immunoblot shows total protein levels of Wpl1-6HA (*α*-HA) with Pgk1 loading control (α-Pgk1). (B) Flow cytometry profiles and (C) nuclear division (n=200) show the timing of bulk DNA replication and chromosome segregation, respectively. (D) Eco1 is essential for meiosis, even in the absence of Wpl1. Spore viability of wild type (AM24170), *wpl1Δ* (AM24265), *eco1-aa* (AM24171), and *eco1-aa wpl1Δ* (AM24289) strains, sporulated in the presence of 1 μM rapamycin. (E and F) Establishment of centromeric cohesion requires Eco1-dependent antagonism of Wpl1. Wild type (AM27183), *wpl1Δ* (AM27186), *eco1-aa* (AM27185), and *eco1-aa wpl1Δ* (AM27184) anchor away strains carrying heterozygous *CEN5*-GFP and *ndt80Δ* were induced to sporulate in 1 μM rapamycin and the percentage of cells with two visible GFP foci was scored at the indicated timepoints (F). Meiotic progression was monitored as DNA content (E). (G and H) Chromosomal arm cohesion requires Eco1, even in the absence of Wpl1. Wild type (AM27253), *wpl1Δ* (AM27256), *eco1-aa* (AM27255), and *eco1-aa wpl1Δ* (AM27254) anchor away strains carrying heterozygous *LYS2*-GFP and *ndt80Δ* were treated and analyzed as described in (E and F). In (E-H) an average of 3 biological replicates is shown; 100 cells were scored for each timepoint in each experiment. Error bars show standard error; * p<0.05, **p<0.01, paired student T test when compared to wild type.

### Distinct requirements for Eco1 in cohesion establishment at centromeres and on chromosome arms

To determine whether *eco1-aa* cells establish functional cohesion during S phase, we labelled one homolog either at a centromere (*CEN5-GFP*) or at a chromosomal arm site (*LYS2*-GFP) and scored the percentage of cells with two GFP foci (indicating defective cohesion) as cells progressed through meiotic S phase into a prophase I arrest. In wild type prophase cells, sister chromatids are tightly cohered and a single focus is visible. In contrast, *eco1-aa* cells showed a profound cohesion defect (Figure 2E-H). Remarkably, deletion of *WPL1* restored cohesion at the centromere (*CEN5-GFP*), but not at the chromosomal arm site (*LYS2-GFP*) (Figure 2F and H). Deletion of *WPL1* alone also caused a modest cohesion defect at *LYS2*, but not at *CEN5* (Figure 2F and H). Therefore, the critical role of Eco1 in cohesion establishment at centromeres is to counteract Wpl1, while on chromosome arms Eco1 plays an additional, essential function.

### Eco1 counteracts Wpl1-dependent cohesin destabilization during meiotic prophase

While Wpl1 promotes cohesin removal from chromosomes between prophase exit and metaphase I (Challa et al., 2019), paradoxically, Wpl1 levels and the slower migrating, presumed active, form decline at prophase exit and are instead highest during meiotic S and prophase (Figure 2A). Therefore, Wpl1 may need to be counteracted by Eco1 prior to or during meiotic prophase. Calibrated ChIP-Seq revealed a global increase in the levels of the cohesin subunit Smc3 on chromosomes of *wpl1Δ* prophase I cells (Figure 3A and B). Meiotic stage and Smc3 total protein levels were comparable in all conditions (Figure 3B and C). In contrast, chromosomal Smc3 was reduced in *eco1-aa* genome-wide, with the notable exception of core centromeres (Figure 3D), Interestingly, inactivation of Wpl1 increased the chromosomal levels of Smc3 in *eco1-aa*, particularly at centromeres, where Smc3 levels in *eco1-aa wpl1Δ* were comparable to the *wpl1Δ* mutant alone (Figure 3D). Elsewhere, including at pericentromere borders and known chromosomal arm cohesin sites (Paldi et al., 2020), Smc3 levels in *eco1-aa wpl1Δ* were similar to wild type (Figure 3D). ChIP-Seq of the meiosis-specific kleisin Rec8 in prophase I revealed a similar pattern to that of Smc3 (Figure S3). Although the different strains cannot be directly compared due lack of a suitable antigen for calibration (see materials and methods), inspection of Rec8 levels in individual strains confirmed that Eco1 is more important for Rec8 association with borders and arm sites than centromeres (Figure S3). Taken together, our Rec8 and Smc3 ChIP-Seq show that the function of Eco1 at pericentromere borders and chromosome arms in meiotic prophase is two-fold. First, Eco1 protects border and arm cohesin from Wpl1-dependent removal, since cohesin levels at these sites are higher in *eco1-aa wpl1Δ* compared to *eco1-aa* cells. Second, Eco1 has an additional, Wpl1-independent function in cohesin retention at border and arm sites, because cohesin levels are higher in *wpl1Δ* compared to *eco1-aa wpl1Δ* cells. In contrast, at centromeres, Wpl1 removes cohesin, but Eco1 has little influence on cohesin levels. This implies that specialized cohesin loading and/or anchoring mechanisms at kinetochores (Hinshaw et al., 2017; Paldi et al., 2020) are the main antagonists of the cohesin removal activity of Wpl1 at centromeres, at least in prophase. Nevertheless, the ability of Wpl1 deletion to rescue the centromeric cohesion detects of *eco1-aa* cells indicates that Eco1 must counteract Wpl1 at centromeres to allow cohesion establishment during S phase (Figure 2F).

**Figure 3.**
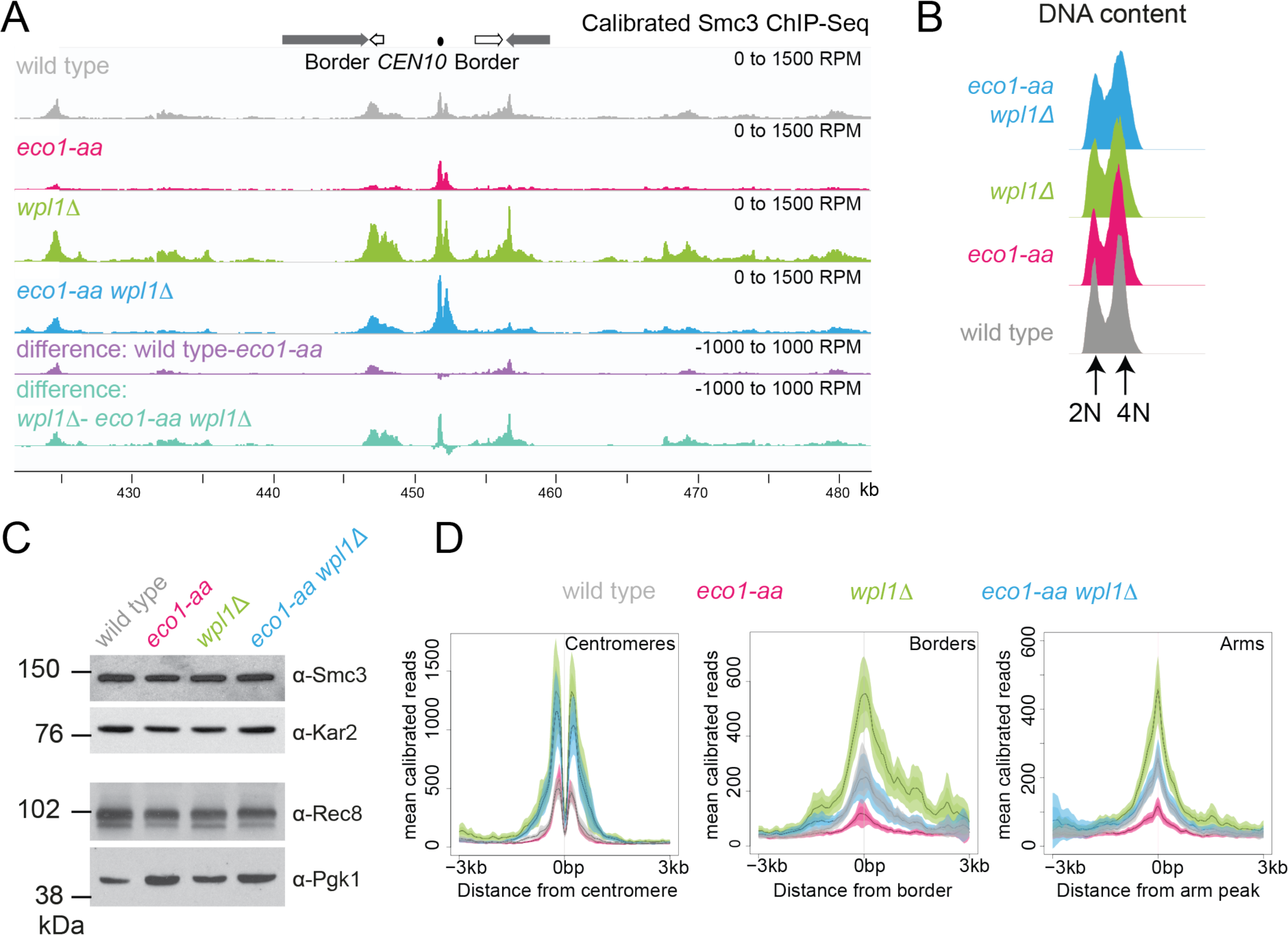
Eco1 restricts Wpl1-dependent removal of chromosomal cohesin during meiotic prophase and retains cohesin at pericentromere borders. Wpl1 globally reduces chromosomal cohesin levels, while Eco1 is required for normal cohesin levels on chromosome arms but not centromeres. Wild type (AM28719), *eco1-aa* (AM28720), *wpl1Δ* (AM29750) and *wpl1Δ eco1-aa* (AM29781) anchor-away strains carrying *ndt80Δ* were harvested 6h after induction of sporulation. (A) Calibrated Smc3 ChIP-seq for a representative region surrounding *CEN10*. (B) Flow cytometry profiles show similar DNA content at harvesting in all cultures. (C) Western immunoblot with loading controls (*α*-Kar2 and *α*-Pgk1)) shows comparable Smc3 (*α*-Smc3) and Rec8 (*α*-Rec8) levels in all cultures at the time of harvesting. (D) Mean calibrated ChIP-Seq reads (line), standard error (dark shading) and 95% confidence interval (light shading) at all 16 centromeres, 32 borders and 32 arm peaks.

### Eco1 establishes loop boundaries in meiotic prophase chromosomes and restricts long-range chromatin interactions independently from Wpl1

Meiotic prophase chromosomes are highly structured, with ordered chromatin loops emanating from linear protein axes which are zipped together by the synaptonemal complex (Figure 4A). To understand how Eco1 and Wpl1 define chromosome structure in meiotic prophase, we performed Hi-C 6h after inducing *ndt80Δ* cells to sporulate and confirmed consistent DNA content for all conditions (Figure 4B). Analysis of contact probability on chromosome arms as a function of genomic distance revealed an increase in long-range interactions in both *eco1-aa* and *wpl1Δ* mutants, which was further exacerbated in *eco1-aa wpl1Δ* cells (Figure 4C). Plotting the slope resulted in a right-ward shift of the curve maxima, which estimates the average size of the loops (Dauban et al., 2020; Figure 4C). This indicated an increase from ∼10kb in wild type to ∼20kb in *eco1-aa* or *wpl1Δ* cells and up to ∼40 kb in *eco1-aa wpl1Δ* (Figure 4C). Heat maps of individual chromosomes corroborated the additive increase in long range interactions in *wpl1Δ eco1-aa* double mutants (Figure 4D). Further inspection revealed that spots and stripes on the Hi-C contact maps, indicative of positioned loops anchored on two or one sides, respectively, were stronger and increased in number in *wpl1*Δ, but were more diffuse in *eco1-aa*, even after Wpl1 deletion (Figure 4D). This suggests loop boundaries/anchors are strengthened by *wpl1Δ* but lost in *eco1-aa.* Mirrored pile-ups of all 16 wild type pericentromeres centered on the centromeres confirmed the organization of the flanking chromatin into two separate domains, indicating that centromeres act as insulators. In contrast, the absence of active Eco1 reduced insulation across centromeres (Figure 4E see also top right and bottom left quadrants on the ratio difference maps in Figure S4E). Interestingly, the length and intensity of the Hi-C contact stripe protruding from centromeres progressively increased in *eco1-aa* and *wpl1Δ eco1-aa* cells, compared to wild type (Figure 4E, arrows; Figure S4E), suggesting that Eco1 limits the extent of loop-extrusion by cohesin complexes anchored at centromeres. Pile-ups centered on all 32 pericentromere borders (Paldi et al., 2020) revealed strong boundaries in wild type, which increased in intensity in *wpl1Δ* cells (Figure 4F; Figure S4F). In contrast, boundaries at pericentromere borders were barely detectable in *eco1-aa* and were only partially rescued by deletion of *WPL1* (Figure 4F), suggesting that Eco1 is critical to halt loop extrusion at borders, even in the absence of Wpl1. Note that border pile-ups also display a second centromere-proximal stripe, corresponding to loop extrusion from the centromeres that is increased in intensity in *wpl1Δ*, consistent with the centromere pile-ups. Both *eco1-aa* and *eco1-aa wpl1Δ* cells also exhibited reduced insulation at borders, indicating that Eco1 is also important to prevent loop extrusion across borders (Figure 4F, see also difference maps in Figure S4F). Consistent with the pile-up analysis, examination of individual wild-type pericentromeres revealed the presence of Hi-C spots, indicative of positioned loops, between the centromere and each of the two borders, as marked by the characteristic tripartite Smc3 ChIP-seq signal (Figure 4G). While stronger Hi-C spots were localized with tripartite Smc3 in *wpl1Δ* cells, both features were absent in *eco1-aa* and weaker in *eco1-aa wpl1Δ* cells (Figure 4G). These data indicate that Wpl1 and Eco1 limit loop expansion through distinct mechanisms in meiotic prophase. While Wpl1 destabilizes loop-extruding cohesin to reduce its lifetime, Eco1 anchors cohesin at boundary sites to stabilize and position loops. We further note that Eco1-dependent loop stabilization critically defines the boundaries that demarcate the pericentromeric domain. This is consistent with the requirement of Eco1 to maintain cohesin association with borders and chromosomal arm sites (Figure 3).

**Figure 4.**
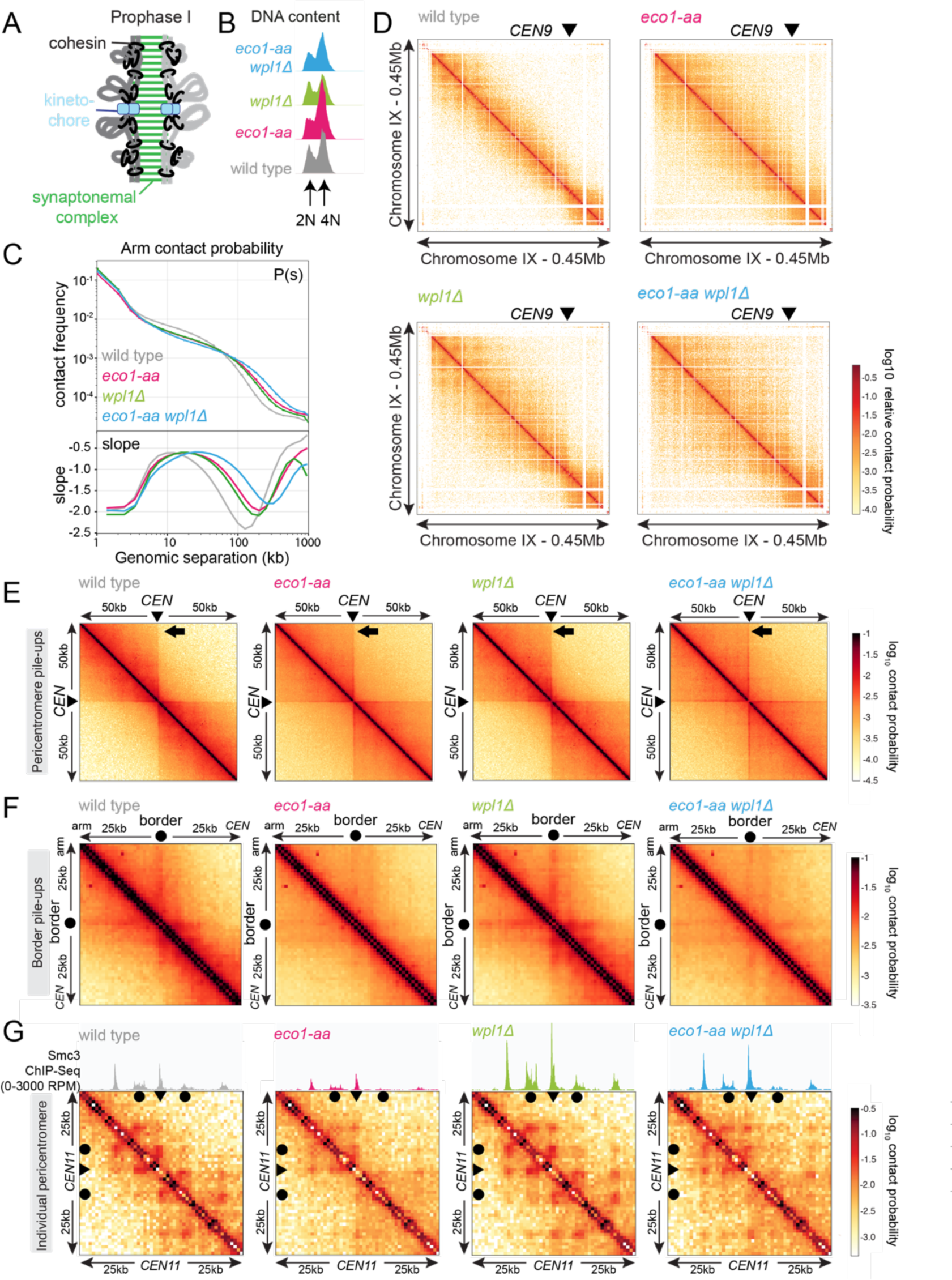
Eco1 restricts long-range chromatin interactions and establishes chromatin boundaries at pericentromere borders. Hi-C analysis of chromosome conformation in meiotic prophase of wild type (AM28719), *eco1-aa* (AM28720), *wpl1Δ* (AM29750) and *eco1-aa wpl1Δ* (AM29781) cells carrying *ndt80Δ*. Strains were harvested 6h after induction of sporulation. (A) Schematic representation of homologous chromosomes emanating from a proteinaceous axis and depicting intra- and inter-sister chromatid cohesin in meiotic prophase. (B) Flow cytometry confirms similar staging in all strains. (C) Contact probability versus genomic distance (*P(s)*) for the indicated strains, excluding contacts across centromeres (1kb bin; log10 scale). The derivative of the P(s) curve (slope) plotted against genomic distance is shown below. (D) Contact maps (1kb bin) show individual chromosome IX for the indicated genotypes. The arrowhead indicates the position of the centromere (*CEN9*). (E) Pile-ups (1kb bin) of *cis* contacts in the 100kb surrounding all 16 centromeres (mirrored). The arrow marks aggregate contacts between the centromeres and the region ∼50kb away on the arm of the same chromosome. (F) Pile-ups (1kb bin) of *cis* contacts in the 50 kb flanking all 32 borders. Pile-ups are oriented so that chromosomal arm and centromere flanks are at the upper left and lower right, respectively. (G) Contact maps (1kb bin) of the pericentromeric region of chromosome XI (50 kb flanking *CEN11*). Calibrated Smc3 ChIP-seq signal for the appropriate genotype (from Figure 3A-D) is shown above. Arrowheads mark the position of centromeres, filled circles mark the borders.

### Replication-independent Smc3 acetylation defines meiotic chromosome loops

Our findings show that Eco1 is a key determinant of loop anchors at pericentromere borders and other chromosomal boundaries in meiotic prophase cells. Interestingly, Eco1-dependent Smc3 acetylation also occurs in unreplicated cells, suggesting that loop anchors may form independently of DNA replication and the presence of a sister chromatid (Figure 1E, Figure S1D). As such, Eco1 could form boundaries by directly acetylating loop extruding cohesin complexes, rather than complexes engaged in cohesion. To test this idea, we generated Hi-C maps of prophase I wild-type and *clb5Δ clb6Δ* cells. Flow cytometry confirmed that, in contrast to wild-type, *clb5Δ clb6Δ* cells failed to undergo bulk pre-meiotic DNA replication (Figure 5A). Although chromosome axis proteins assemble apparently normally in *clb5Δ clb6Δ* cells, double strand break formation and, consequently, homolog pairing are defective (Blitzblau et al., 2007; Brar et al., 2009; Smith et al., 2001). Therefore, both inter-sister and inter-homolog interactions are expected to be absent in *clb5Δ clb6Δ* cells, so that any stripes and dots observed on Hi-C maps can be attributed to *cis-* looping along a single chromatid. Consistently, in *clb5Δ clb6Δ* cells, mid- to long-range contacts on individual chromosomes were strongly reduced, though the characteristic stripe and dot pattern indicating the presence of loops was still visible (Figure 5B). Analysis of contact probability on chromosome arms found that contacts in the 5-100kb range are reduced in *clb5Δ clb6Δ* cells (Figure 5C). Importantly, however, plotting the derivative revealed that the average loop size in *clb5Δ clb6Δ* was only slightly reduced (∼8kb in *clb5Δ clb6Δ* vs 10kb in wild type; Figure 5C). This suggests that the decrease in mid-/long-range interactions might result from the absence of inter-sister and inter-homolog contacts. Together with the observation that *clb5Δ clb6Δ* cells do not display an increase in long range interactions, this indicates that replication and the presence of cohesion do not play a fundamental role in restricting loop extrusion. To confirm the notion that loop boundaries form in unreplicated cells, we examined individual pericentromeres of *clb5Δ clb6Δ* cells. As an example, Figure 5D shows that Hi-C spots and stripes are anchored at the pericentromere borders of chromosome IX in wild type and *clb5Δ clb6Δ* cells, respectively. The presence of Hi-C stripes, rather than spots, in *clb5Δ clb6Δ* cells suggests that *cis*-looping may be more dynamic than in replicated cells, perhaps due to the absence of sister chromatid cohesion. Centromere and border Hi-C pile-ups for the wild type and *clb5Δ clb6Δ* Hi-C datasets further corroborated these findings (Figure 5E-H). Interestingly, centromere pile-ups and ratio maps revealed that the contact stripe originating from centromeres was greatly diminished in *clb5Δ clb6Δ* and accompanied by a loss of centromeric insulation (Figure 5E and F). Although the underlying reasons are unclear, this suggests that Clb5 and Clb6 promote centromere-directed loop extrusion. In contrast, border pile-ups confirmed the presence of boundaries in *clb5Δ clb6Δ* cells, though they were not as strong as in wild type (Figure 5G and H). Overall, these results indicate that although replication is not required for boundary formation, cohesion and/or the presence of a sister chromatid strengthen these boundaries. Importantly, boundary formation and loop anchoring by pericentromere borders in *clb5Δ clb6Δ* contrasts with *eco1-aa* cells which show greatly diminished border-anchored loops (compare Figure 5D with Figure 4G and Figure 5H with Figure S4B) and border/centromere contacts (compare Figure 5G with Figure 4F) both on individual pericentromeres or on border pile-ups and pile-up ratio plots. Since robust boundary formation in replicated cells relies on Eco1 (Figure 4), and because Eco1 is proficient in acetylating cohesin also in unreplicated cells (Figure 1E and Figure S1D), it follows that Eco1 is capable of anchoring loops independently of DNA replication and sister chromatid cohesion. Moreover, it implies that loop extruding cohesin can be acetylated by Eco1.

**Figure 5.**
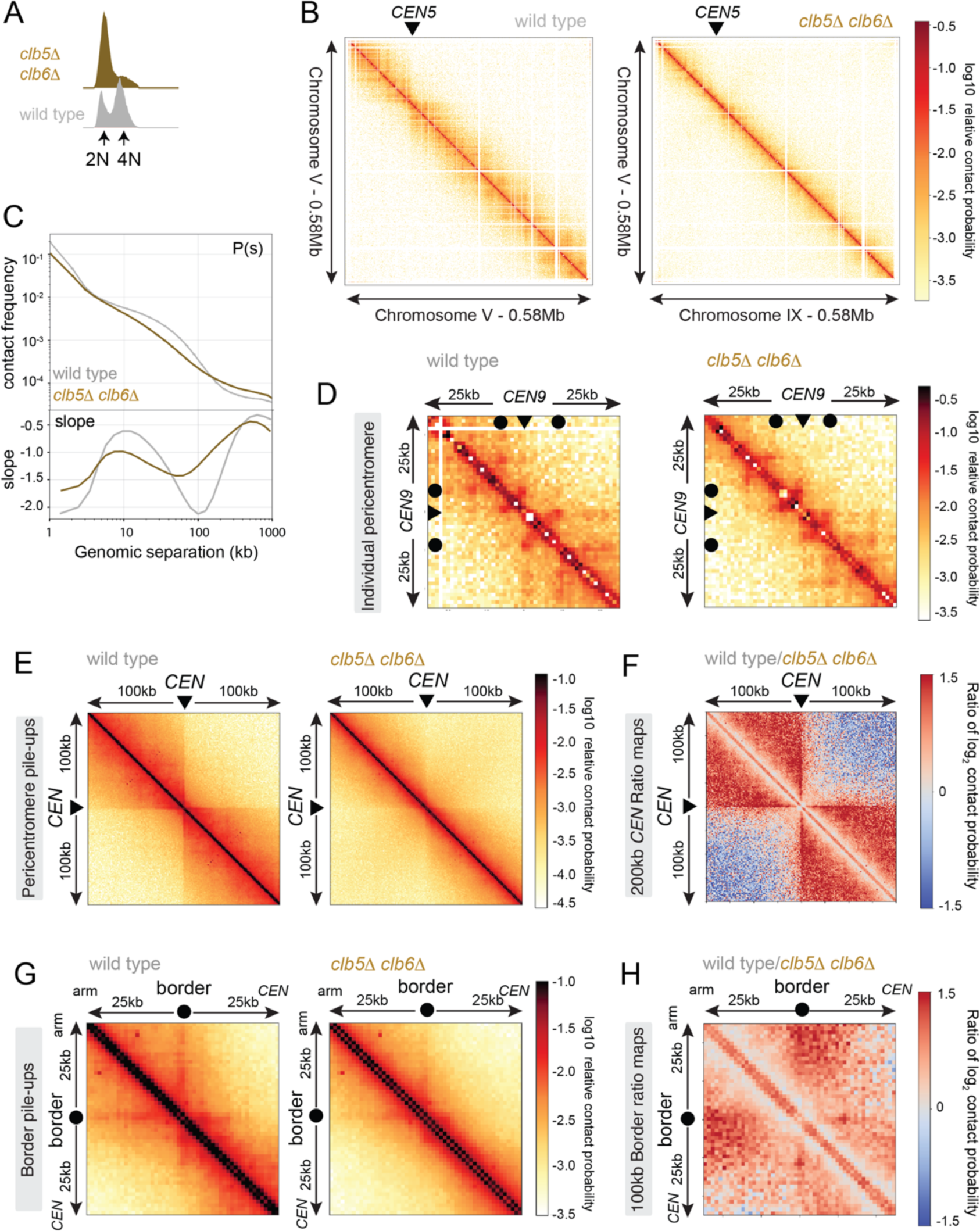
Boundary formation occurs on unreplicated chromosomes. Hi-C analysis of chromosome conformation in meiotic prophase in the absence of DNA replication. Wild type (AM11633) and *clb5Δ clb6Δ* (AM28841) cells carrying *ndt80Δ* were analyzed as described in Figure 4. (A) Flow cytometry profiles confirm that *clb5Δ clb6Δ* fail to undergo meiotic DNA replication. (B) Contact maps (1kb bin) show decreased mid-to-long range contacts in *clb5Δ clb6Δ* on individual chromosome V. The arrowhead indicates the position of the centromere (*CEN5*). (C) Contact probability versus genomic distance (*P(s)*), excluding contacts that occur across centromeres (1kb bin; log10 scale). The derivative of the P(s) curve (slope) plotted against genomic distance is shown below. (D) Contact maps (1kb bin) of the pericentromeric region of chromosome IX (50 kb flanking *CEN9*). Arrowheads mark the position of centromeres, filled circles mark the borders. (E) Pile-ups (1kb bin) of *cis* contacts in the 100kb surrounding all 16 centromeres (mirrored). (F) Ratio plots of *CEN* pile-ups in (E). (G) Pile-ups (1kb bin) of *cis* contacts in the 50 kb flanking all 32 borders. Pile-ups are oriented so that chromosomal arm and centromere flanks are at the upper left and lower right, respectively. (H) Log2Ratio plots of border pile-ups in (G).

### Recombination prevents prophase exit in eco1-aa cells

We next sought to understand how boundary formation and cohesion establishment by Eco1 impact meiotic chromosome segregation. Though *eco1-aa* cells complete bulk DNA replication in meiotic S phase with similar timing to wild type (Figure 2E and G), only a small fraction of cells undergo nuclear divisions (Figure 1I). Cohesin is required for meiotic recombination and *rec8Δ* cells undergo a recombination-dependent checkpoint arrest in meiotic prophase, due to the persistence of unrepaired double strand breaks (Klein et al., 1999). We found that *eco1-aa* cells similarly arrest in prophase as judged by a failure to separate spindle pole bodies (SPBs, marked by Spc42-tdTomato) and prophase exit was only modestly advanced by deletion of *WPL1* (compare *eco1-aa* to *wpl1Δ eco1-aa* cells; Figure 6A). This indicates that although Eco1 facilitates prophase exit by counteracting Wpl1, other Eco1 functions are also important. To determine whether activation of the recombination checkpoint prevents timely prophase exit in *eco1-aa* and *eco1-aa wpl1Δ* cells, we abolished meiotic double strand break formation (by deletion of *SPO11*). Figure 6A shows that *SPO11* deletion abolished the prophase exit delay of both *eco1-aa* and *eco1-aa wpl1Δ* cells, confirming that Eco1 is required for satisfaction of the recombination checkpoint to allow prophase exit.

**Figure 6.**
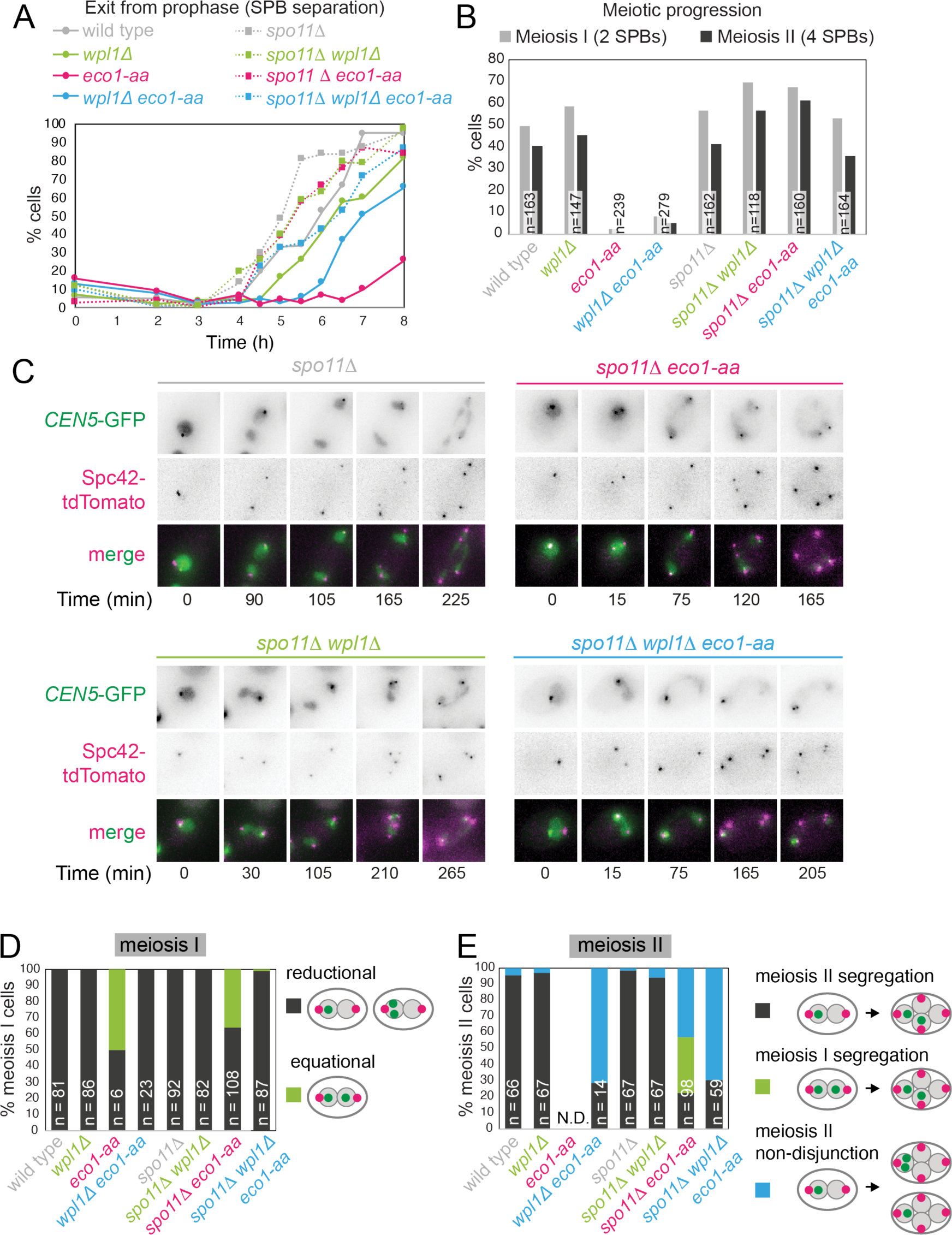
Critical Wpl1-dependent and independent roles of Eco1 in chromosome segregation during both meiosis I and II. (A) Meiotic double strand break formation arrests *eco1-aa* cells in meiotic prophase. Wild type (AM24167), *wpl1Δ* (AM24168), *eco1-aa* (AM24184), *eco1-aa wpl1Δ* (AM24169), *spo11Δ* (AM27670) *spo11Δ wpl1Δ* (AM27673), *spo11Δ eco1-aa* (AM27672) and *spo11Δ eco1-aa wpl1Δ* (AM27671) anchor away strains carrying heterozygous *SPC42-tdTOMATO* and *CENV*-GFP were induced to sporulate. The percentage of cells with more than one Spc42-tdTomato foci was determined from cell populations fixed at the indicated timepoints after resuspension in SPO medium containing rapamycin. At least 100 cells were scored per timepoint. (B-E) Live cell imaging of strains as in (A) sporulated in the presence of rapamycin. (B) Deletion of *SPO11* restores meiotic progression to *eco1-aa* cells. Completion of meiosis I (two Spc42-tdTomato foci) and II (four Spc42-tdTomato foci) was scored. (C) Representative images of *spo11Δ* background cells undergoing the meiotic divisions. (D) Eco1 counteracts Wpl1 to allow sister kinetochore monoorientation. Segregation of *CEN5*-GFP foci to the same (reductional; dark grey) or opposite (equational; green) poles was scored in meiosis I (as two Spc42-tdTomato foci separate; binucleate cells). (E) Eco1 is required for pericentromeric cohesion, even in the absence of Wpl1. Segregation of *CEN5*-GFP foci to opposite (dark grey) or the same pole(s) (blue) was scored in during meiosis II (as four Spc42-tdTomato foci separate; tetranucleate cells). Cells that had already segregated their sister *CEN5*-GFPs in meiosis I (GFP foci in two nuclei at the binucleate stage) were scored as a separate category (green).

### Co-segregation of sister chromatids during meiosis I requires Wpl1 antagonism by Eco1

To understand how Eco1/Wpl1-dependent chromosome organization impacts meiotic chromosome segregation, we exploited the ability of *spo11Δ* to bypass of the prophase block of *eco1-aa* cells. Note that Eco1-dependent cohesin acetylation does not require recombination (Figure S1E). Imaging of live cells carrying a heterozygous centromere label (*CEN5-*GFP) and Spc42-tdTomato confirmed that deletion of *SPO11* permitted meiotic divisions in *eco1-aa* and *eco1-aa wpl1Δ* (Figure 6B). In wild type cells, sister chromatids co-segregate in meiosis I (“reductional segregation”) so that *CEN5-*GFP foci are inherited by just one of the two nuclei and this was also the case in *spo11Δ*, *wpl1Δ* and *spo11Δ wpl1Δ* cells, (2 separated SPBs; Figure 6C and D; Figure S5A). However, surprisingly, in ∼40% of *spo11Δ eco1-aa* cells, *CEN5-*GFP foci segregated to opposite poles in meiosis I (“equational segregation”) indicating loss of sister kinetochore monoorientation (Figure 6C and D). Remarkably, deletion of *WPL1* largely rescued the sister kinetochore monoorientation defect of *spo11Δ eco1-aa* cells in meiosis I (Figure 6C and D), consistent with the restoration of centromere cohesion at prophase in *eco1-aa wpl1Δ* compared to *eco1-aa* cells (Figure 2F). This supports the idea that Eco1 may enable sister kinetochore monoorientation through effects on cohesion establishment. Therefore, Eco1 antagonism of Wpl1 allows the establishment of centromeric cohesion and enables sister kinetochore monoorientation.

### Sister chromatid segregation during meiosis II requires an Eco1 function distinct from Wpl1 antagonism

During wild type meiosis II, segregation of sister chromatids to opposite poles is ensured by the pool of cohesin that persists on pericentromeres after bulk cohesin cleavage in anaphase I and which likely resides at borders. In wild type cells, this can be visualized by sister *CEN5-*GFP foci segregation in anaphase II, resulting in their association with two of the four Spc42-tdTomato foci (Figure S5A). Similarly, *spo11Δ, wpl1Δ*, and *spo11Δ wpl1Δ* cells segregated sister *CEN5-GFP* foci to opposite poles in meiosis II (Figure 6C and E; Figure S5A). In *spo11Δ eco1-aa* cells, among the cells that enter meiosis II ∼30% had already segregated sister *CEN5-* GFP foci during meiosis I (green bar in Figure 6E), while in a further ∼50% of cells, *CEN5-*GFP were found next to a single Spc42-tdTomato focus (blue bar in Figure 6E), indicating defective meiosis II segregation. Moreover, defective meiosis II segregation in *eco1-aa* cells is not rescued by deletion of *WPL1*, whether or not Spo11 is present (Figure 6E). Therefore, Eco1 is essential for chromosome segregation during meiosis II, even in the absence of Wpl1. One potential explanation for these findings is that localization of the cohesin protector protein, shugoshin (Sgo1) or the meiotic protein Spo13, which is also required for cohesin protection during meiosis I (Galander et al., 2019b, 2019a; Katis et al., 2004; Lee et al., 2004), may require Smc3 acetylation. However, ChIP-qPCR revealed that both Sgo1 and Spo13 localize to chromosomes in *eco1-*aa cells (Figure S5B-F). Both proteins follow a similar pattern to Rec8, showing reduced chromosomal association in *eco1-aa* cells that is rescued by *WPL1* deletion (Figure S5B-F), consistent with a requirement for cohesin for the chromosomal association of Sgo1 and Spo13 (Galander et al., 2019b; Kiburz et al., 2005). Therefore, the failure to build, rather than protect, pericentromeric cohesion is the cause of meiosis II mis-segregation in *eco1-aa* cells. Since Wpl1 rescues the loss of cohesin on pericentromeric borders in *eco1-aa* cells, but not the anchoring of loops, it is likely that cohesin anchoring at chromatin boundaries is the critical function of Eco1 in cohesion establishment and meiosis II segregation.

### Mutation of Smc3-K112,113 results in meiotic lethality

In mitotically-growing cells, Eco1 protects cohesin from Wpl1 by acetylation of Smc3 residues K112 and K113 (Ben-Shahar et al., 2008; Rowland et al., 2009; Unal et al., 2008). To determine whether Smc3-Ac similarly allows cohesion establishment by protecting from Wpl1 and/or confers chromatin boundary function in meiosis, we analyzed a non-acetylatable *smc3-K112,113R* mutant. To support mitotic growth, cells carried wild type *SMC3* under the *CLB2* promoter, which is repressed in meiosis (homozygous *pCLB2-3HA-SMC3*), and heterozygous *smc3-K112,113R* or, as a control *SMC3,* at an ectopic locus expressed from the endogenous promoter (Figure S6A). Smc3 levels in *pCLB2-3HA-SMC3* cells without ectopic expression were largely repressed, though low levels of residual Smc3 were detectable (Figure S6B and C). Levels of heterozygously produced Smc3 and Smc3-K112,113R were approximately half that of Smc3 in wild type cells (Figure S6B and C). Ectopic *SMC3* expression rescued the unviability of *pCLB2-3HA-SMC3* spores while *smc3-K112,113R* expression did not, indicating that Smc3 acetylation is essential for meiosis (Figure S6D). To determine whether Smc3-K112,113R localizes normally to chromosomes and whether it is susceptible to destabilization by Wpl1, we performed calibrated Smc3 ChIP-Seq in prophase I cells where either Smc3 or Smc3-K112,113R were heterozgously produced and in the presence and absence of Wpl1 (Figure 7A-D). Flow cytometry confirmed similar meiotic progression (Figure 7A) and western blotting showed that total cellular Smc3 levels were comparable (Figure 7B). Like *eco1-aa* (Figure 3A-D), *smc3-K112,113R* reduced chromosomal cohesin levels genome-wide, except at centromeres. *WPL1* deletion increased Smc3 levels in *smc3-K112,113R* cells (Figure 6C), but to a lesser extent than in the *eco1-aa* background (Figure 3A). Although the reasons for this are unclear, the amino acid substitutions in Smc3 may themselves perturb cohesin loading or translocation, in addition to preventing acetylation by Eco1. Nevertheless, although introduction of the *smc3-K112,113R* mutation into *wpl1Δ* cells only slightly reduced Smc3 levels at centromeres, *smc3-K113,113R* greatly reduced Smc3 levels at pericentromeric borders and arm sites (Figure 7C-D, compare *wpl1Δ* and *smc3-K112,113R wpl1Δ*). Therefore, like Eco1, Smc3-Ac is more important for retention of cohesin at pericentromere borders and arm sites than at centromeres, suggesting a role in boundary formation.

**Figure 7.**
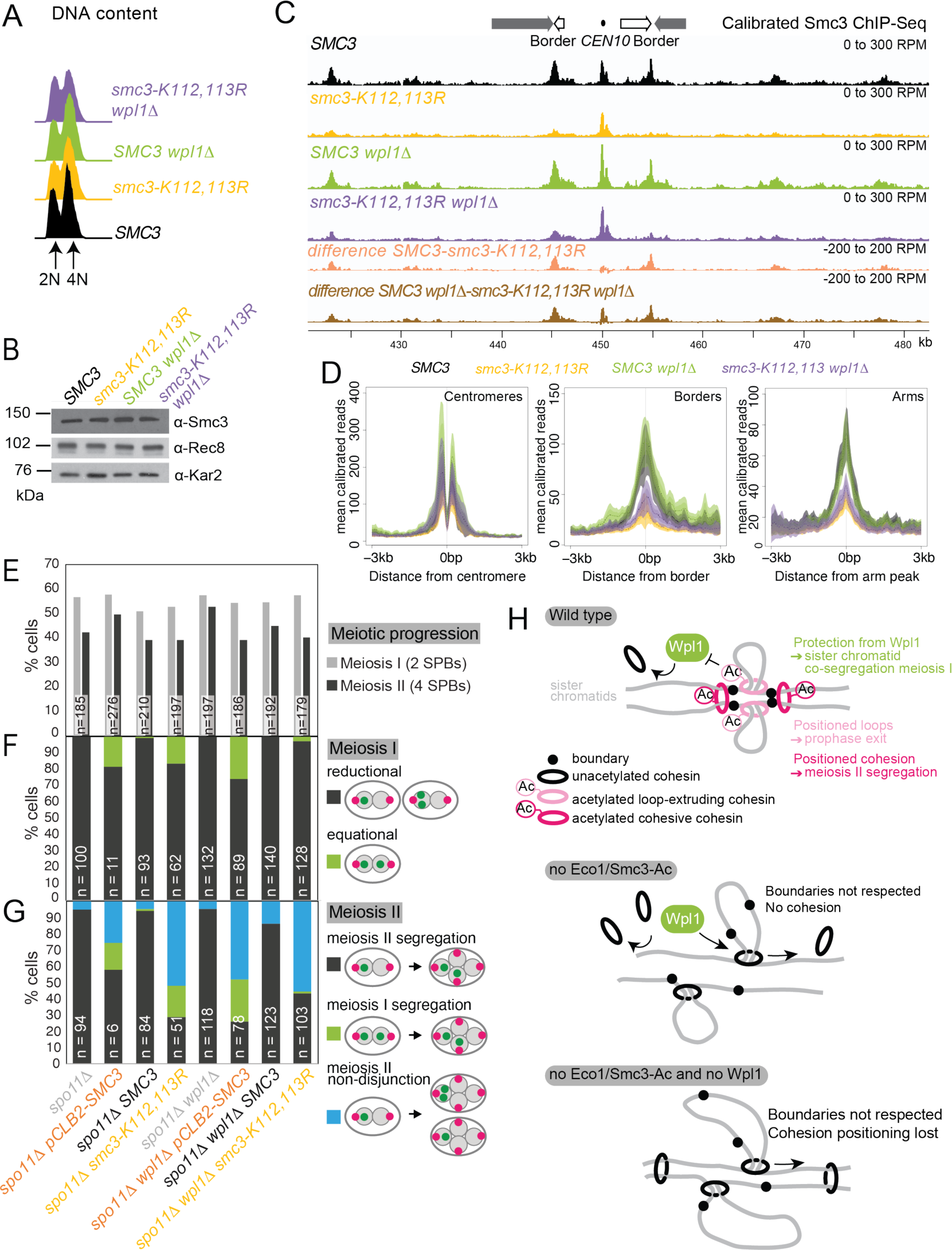
Smc3 acetylation is essential for meiosis. (A-D) *smc3-K112,113R* leads to a global reduction in chromosomal Smc3 levels, which is only partially restored by *WPL1* deletion. *SMC3* (AM29315), *smc3-K112,113R* (AM29316), *SMC3 wpl1Δ* (AM30310) and *smc3-K112,113R wpl1Δ* (AM30311) strains carrying *ndt80Δ* were harvested 6h after induction of sporulation. (A) Flow cytometry profiles show similar DNA content at harvesting in all cultures. (B) Western immunoblot with Kar2 loading control (*α*-Kar2) shows comparable Smc3 (*α*-Smc3) and Rec8 (*α*-Rec8) levels in all cultures at the time of harvesting. (C) Calibrated Smc3 ChIP-seq for a representative region surrounding *CEN10*. (D) Mean calibrated ChIP-Seq reads (line), standard error (dark shading) and 95% confidence interval (light shading) at all 16 centromeres, 32 borders and 32 flanking arm sites. (E-G) Smc3 acetylation is required to ensure co-segregation of sister chromatids in meiosis I and accurate meiosis II chromosome segregation. Meiotic progression (E) and meiosis I (F) and II (G) chromosome segregation were scored after live cell imaging as in Figure 6 (B-E). Strains used were *spo11Δ* (AM30238), *spo11Δ pCLB2-3HA-SMC3* (AM30240), *spo11Δ SMC3* (AM30242), *spo11 smc3-K112,113R* (AM30244), *spo11Δ wpl1Δ* (AM30234), *spo11Δ wpl1Δ pCLB2-3HA-SMC3* (AM30235), *spo11Δ wpl1Δ SMC3* (AM30655) and *spo11Δ wpl1Δ smc3-K112,113R* (AM30237). (H) Model for Eco1 and Wpl1 roles in meiosis. In wild type cells (top panel), Eco1 cohesin acetylation is essential in three meiotic processes: it protects centromeric cohesin from Wpl1-mediated release, allowing sufficient cohesion to be built to establish monoorientation; it positions DNA loops along chromosome arms to promote recombination and prophase exit; and it positions loops and cohesion at pericentromeric borders to guide correct sister chromatid segregation in meiosis II. In the absence of Eco1 or Smc3-Ac (middle panel), boundaries are not respected and more cohesin complexes are released from DNA due to the action of Wpl1, with detrimental effects on recombination and meiosis I and II segregation. In the absence of both Eco1 or Smc3-Ac and Wpl1, cohesion is partially restored, specifically at the centromere, but loop boundaries are not, leading to the formation of long unpositioned loops that are not able to support prophase recombination and meiosis II segregation.

Next we asked whether Smc3-Ac underlies the functional effects of Eco1 in meiosis by conferring both cohesion establishment and boundary function. First, we scored sister *CEN5-*GFP or *LYS2-*GFP foci separation as cells progressed into a prophase arrest and found that *smc3-K112,113R* cells exhibited similar cohesion defects to cells depleted of Smc3 (Figures S6E-J). In both *smc3-K112,113R* and *pCLB2-SMC3* cells, the cohesion defect at centromeres was more modest than that of *eco1-aa* cells (Figure 2F), potentially reflecting incomplete meiotic depletion of Smc3 when placed under the *CLB2* promoter (Figure S6B and C). Next, we used live-cell imaging to assay the requirement for Smc3-Ac in prophase exit and meiosis I and II chromosome segregation. Exit from prophase was impaired in *pCLB2-3HA-SMC3* and, to a lesser extent, *smc3-K112,113R* cells, but in both cases was overcome by deletion of *SPO11* (Figure S7A; Figure 7E), indicating that Smc3-Ac is required for satisfaction of the recombination checkpoint, like Eco1. Analysis of sister chromatid segregation in meiosis I in the *spo11Δ* background (Figure 7F; Figure S7B) revealed that ∼10-15% of *pCLB2-SMC3* and *smc3-K112, 113R* cells aberrantly segregated sister chromatids to opposite poles during meiosis I (Figure 7F, Figure S7B). In the case of *smc3-K112,113R*, but not *pCLB2-SMC3,* this equational meiosis I segregation was rescued by deletion of *WPL1* (Figure 7F). Sister chromatid segregation during meiosis II was also greatly impaired in both *pCLB2-SMC3* and *smc3-K112,113R* cells, but in neither case was it rescued by *WPL1* deletion (Figure 7G and Figure S7C). We note that *smc3-K112,113R* and *pCLB2-SMC3* have a lesser effect on meiosis I sister chromatid segregation and centromeric cohesion as compared to *eco1-aa* (compare Figure 2F with Figure S6E). In contrast, meiosis II segregation was similarly defective in *eco1-aa, pCLB2-SMC3* and *smc3-K112,113R* cells (Figure 6E and Figure 7G). The most likely explanation for this difference is that pre-meiotic expression of *pCLB2-SMC3* leads to persistence of functional Smc3 (Figure S6C) which is preferentially loaded at centromeres to partially establish cohesion at this site. However, in both meiosis I and II segregation *WPL1* deletion had the same effect on *eco1-aa* and *smc3-K112,113R* cells. We conclude that Eco1-dependent Smc3-Ac functionally organizes meiotic chromosomes for their segregation. At centromeres, Eco1-dependent Smc3 acetylation counteracts Wpl1 to direct sister kinetochore monoorientation and thereby enforce sister chromatid co-segregation in meiosis I. Eco1 acetylation of Smc3 is also essential for prophase exit and sister chromatid segregation in meiosis II, even in the absence of Wpl1, likely as a result of cohesin anchoring to establish chromatin boundaries.

## Discussion

### Cohesin regulators organize functionally distinct chromosomal domains

We have shown how cohesin regulators remodel chromosomes into functionally distinct domains to allow for the unique events of meiosis. We find that the control of loop formation and cohesion establishment by Eco1 and Wpl1 allows centromeres, pericentromeres and chromosome arms to adopt region-specific functions in sister kinetochore co-orientation, cohesion maintenance and recombination. These functions of Eco1 and Wpl1 that we uncovered in meiosis may also explain how other chromosome domains are established to support other genomic functions, including localized DNA repair and control of gene expression.

### Loop anchoring allows the formation of specific chromosomal boundaries

Eco1 associates with replication factors and is proposed to couple cohesion establishment to DNA replication by travelling with replication forks (Ivanov et al., 2018; Ladurner et al., 2016; Lengronne et al., 2006; Song et al., 2012). In mitotically dividing yeast cells, Smc3 acetylation is largely dependent on DNA replication (Ben-Shahar et al., 2008), however during meiotic S phase substantial Smc3 acetylation, and anchoring of chromatin boundaries, occurs even in the absence of DNA replication. Cohesin acetylation also occurs without DNA replication in mammalian G1 cells, with a preference for STAG1-containing cohesin complexes (Alomer et al., 2017; Wutz et al., 2020). Therefore, Smc3-Ac can exist independently of cohesion establishment. Indeed, Smc3-Ac *per se* is not critical for cohesion since mitotic *eco1Δ wpl1Δ* yeast cells build sufficient cohesion to support viability. How Eco1 gains access to cohesin that is not associated with replication forks remains to be understood but it is interesting to speculate that specialized cohesin subunits, for example STAG1 in mammals or Rec8 in yeast, may allow cohesin targeting by Eco1 independent of the replication machinery.

In addition to Wpl1 antagonism, Eco1 and Smc3-Ac confer a more fundamental role in chromosomal loop positioning that we find to be indispensable for meiosis. We envisage that early in meiosis, prior to DNA replication, Scc2-Scc4-cohesin complexes load onto chromosomes and begin to extrude loops (Figure 7H). Loaded cohesin, and loops, have a limited lifetime due to the removal of unacetylated cohesin by Wpl1. However, a fraction of cohesin engaged in *cis-*looping is acetylated by Eco1, with two consequences. First, Smc3-Ac blocks cohesin’s Wpl1-dependent release. Second, Smc3-Ac anchors cohesin at the base of loops, preventing loop migration and restricting extrusion, resulting in the stable positioning of moderately-sized loops. The absence of Wpl1 increases the lifetime of loop-extruding cohesin complexes resulting in loop extension (Figure 4; (Costantino et al., 2020; Dauban et al., 2020; Haarhuis et al., 2017)). The combined absence of both Eco1 and Wpl1 results in mobile cohesin with extended lifetimes on chromosomes and untempered loop extruding activity, leading to long, poorly positioned loops. The positioning activity of Eco1 also facilitates cohesion establishment along chromosome arms by anchoring cohesin complexes that are engaged in linking the two sister chromatids (“cohesive cohesin’). Whether it does this directly by acetylating cohesive cohesin to prevent its migration, or indirectly by generating *cis-*loops that form a barrier to translocation of cohesive cohesin is unclear. At centromeres, Smc3-Ac, which blocks the destabilizing activity of Wpl1, is sufficient for cohesion establishment, likely due to enhanced cohesin loading or specialized anchoring mechanisms at this site. A recent report indicates that Eco1-dependent Smc3 acetylation also establishes loop positioning in S phase of yeast mitotic growth (Bastié et al., 2021). Exactly how acetylation affects cohesin enzymology awaits detailed biochemical analysis.

### Establishment of functional chromosomal units for meiotic recombination

Our studies on meiosis, where chromosomal domains must be defined to lose cohesin at chromosome arms in meiosis I and pericentromeres in meiosis II, provide a unique opportunity to dissect the functional importance of loop positioning. We found that Eco1 and Smc3-Ac are critical for the completion of DNA recombination to allow exit from meiotic prophase. Restoration of cohesin levels in Eco1-deficient cells by removal of Wpl1 was insufficient to support DNA repair and prophase exit, or for chromosomal arm cohesion. We speculate that loss of positioned chromatin loops causes defects in the repair of double strand breaks leading to a delay in prophase exit. Eco1-dependent cohesin anchoring may promote homology search, be required to reel broken ends within loops to chromosome axes for repair and/or ensure that loops are positioned in register with the homologous chromosome. Although positioned loops are readily detectable during meiotic prophase in budding yeast ((Schalbetter et al., 2019) Figure 4), this is not the case in mouse, rhesus monkey, human and mouse spermatocytes (Alavattam et al., 2019; Patel et al., 2019; Wang et al., 2019). Whether this underlies biological differences in mammals or a technical limitation due to the larger genome size remains unclear.

Eco1 is present beyond S phase into prophase, suggesting that it may play an active role in loop repositioning and/or the anchoring of new loops during homologous recombination. However, Eco1 is degraded at prophase exit, after which only existing loop anchors and positioned cohesion will persist. Given the critical importance of cohesion/loop positioning for chromosome segregation (see below), this has important implications for meiosis in human eggs where the events of prophase and the meiotic divisions are temporally separated by years. It will be interesting to understand whether Esco1/2 remain active in mammalian eggs and whether this is important to safeguard cohesin during the female reproductive lifespan.

### Centromeric cohesion directs sister chromatid co-segregation during meiosis I

Centromeres are unique in retaining higher cohesin levels in the absence of Eco1 function, but this is insufficient for centromeric cohesion. Moreover, Wpl1 inactivation in *eco1-aa* cells leads to an increase in chromosomal cohesin genome-wide, but cohesion is rescued specifically at centromeres. Our live-cell imaging revealed that the loss of centromeric cohesion in *eco1-aa* cells is accompanied by the aberrant segregation of sister chromatids to opposite poles in meiosis I, which was also rescued by Wpl1 inactivation. In Smc3-depleted and *smc3-K112,113R* cells, likely due to residual *SMC3* expression from the *CLB2* promoter, the centromeric cohesion defect was more modest and segregation of sister chromatids to opposite poles was less frequent, but was nevertheless rescued by Wpl1 inactivation. Although we cannot currently rule out the possibility that Eco1 has an additional substrate that confers its function in sister kinetochore co-orientation as has been suggested for fission yeast (Kagami et al., 2017), the simplest explanation is that acetylation of Smc3 by Eco1 counteracts Wpl1 at centromeres allowing the establishment of centromeric cohesion, which in turn facilitates kinetochore fusion via the monopolin complex (reviewed in (Duro and Marston, 2015)). Interestingly, budding yeast lacking Rec8 cohesin do not show co-orientation defects (Monje-Casas et al., 2007; Toth et al., 2000), raising the possibility that mitosis-like, Scc1-containing cohesin complexes confer the co-orientation function of cohesin at centromeres. It will be of great interest to understand how monopolin and cohesin-mediated co-orientation pathways intersect to ensure proper meiosis I chromosome segregation.

### Pericentromere boundaries define persistent cohesion at meiosis II

During meiosis, cohesion on chromosome arms requires Eco1 and Smc3-Ac even in the absence of Wpl1. This argues against the idea that the key function of Smc3-Ac in cohesion is to antagonize Wpl1. Instead, we propose that the ability of Eco1 to reinforce chromatin boundaries is critical for cohesion during meiosis. This phenomenon is most apparent at pericentromere borders, where centromere-loaded cohesin is trapped by convergent genes in mitosis (Paldi et al., 2020). In meiosis II, cells rely entirely on cohesin at pericentromere borders to hold sister chromatids together. We now show that Eco1 is required to retain cohesin at these sites and, consequently, to establish the boundaries that structure pericentromeres during meiosis. As a result, sister chromatids undergo random meiosis II segregation in cells lacking Eco1, whether or not Wpl1 is present. This can explain why cohesin anchoring is essential in meiosis but not in mitosis, where cohesin is present also along chromosome arms. Understanding how Eco1-dependent Smc3 acetylation and convergent genes together trap cohesin to elicit loop anchoring and cohesion at pericentromere borders is an important question for the future.

### Importance of cohesin anchoring

A universal feature of cohesin-dependent chromosome organization is the existence of boundary elements which position both loops and cohesion. However, how these boundaries are established and the functional importance of cohesin anchoring were unclear. We found that Eco1-dependent cohesin acetylation confers boundary recognition and cohesin anchoring. The importance of loop anchoring was revealed by their requirement for proper recombination, centromere cohesion and pericentromeric cohesion in meiotic chromosome segregation. Our demonstration of the importance of cohesin anchoring for fundamental chromosomal processes provides a framework for understanding how chromatin folding into defined domains maintains and propagates the genome.

## Acknowledgements

We are grateful to Andreas Hochwagen for yeast strains, Jean-Paul Javerzat, Lori Koch, Flora Paldi and Gerard Pieper for comments on the manuscript, and to Akira Shinohara for helpful discussions. We thank Flora Paldi for advice on Hi-C sample preparation, and Weronika Borek, Julie Blyth and Karolina Lesniewska for technical assistance with antibody generation. We gratefully acknowledge the EMBL Genecore for DNA sequencing, the Wellcome Centre for Cell Biology Core Bioinformatics Facility for sequencing analysis and the Wellcome Centre Optical Imaging Laboratory (COIL) and Dave Kelly for microscopy and flow cytometry support. This work was funded through a Wellcome Senior Research Fellowship [107827] (ALM, LFM, REB), a Wellcome Investigator award [220780] (ALM, LFM), BBSRC grant [BB/S018018/1] (ALM, LFM), a Wellcome PhD studentship [102316] (REB) and core funding for the Wellcome Centre for Cell Biology [203149] (REB, LFM, DR and ALM).

## Author contributions

Conceptualization: ALM, REB and LFM; Methodology: REB, LFM and ALM; Formal Analysis: DR; Investigation: LFM, REB and ALM; Writing – original draft: ALM; Writing – Reviewing and Editing: ALM, LFM, REB and DR. Visualization: ALM, LFM, REB and DR; Supervision: ALM; Funding acquisition: ALM.

## Materials and Methods

### Yeast strains and plasmids

Yeast strains used in this study were either *Saccharomyces cerevisiae* SK1 or W303 derivatives or *Schizosaccharomyces pombe*, all of which are listed in Table S1. Plasmids used in this study are listed in Table S2. Gene tags, gene deletions, and promoter replacements were introduced using standard PCR-based methods. Specific depletion of proteins (Cdc20, Smc3, Cdc6) during meiosis was achieved by placement of genes under the mitosis-specific *CLB2* promoter (Lee and Amon, 2003). For early meiotic block/release experiments strains carried *pCUP1-IME1 pCUP1-IME4* (Berchowitz et al., 2013), for prophase block/release experiments strains carried *pGAL1-NDT80 pGPD1-GAL4(848).* (Benjamin et al., 2003; Carlile and Amon, 2008). For experiments undertaken in a prophase I arrest the strains carried *ndt80Δ* (Xu et al., 1995), while for experiments in a metaphase I arrest cells carried *pCLB2-CDC20*. All anchor-away strains and their controls carried *RPL13A-2xFKBP12 fpr1Δ tor1-1* (Haruki et al., 2008) (kind gift from Andreas Hochwagen (NYU, New York)). For centromeric GFP foci assays strains carried *CEN5::tetOx224 pURA3-TetR-GFP* (Tanaka et al., 2000) and for arm GFP foci assays strains carried *lys2::tetOx240 pURA3-TetR-GFP* (Brar et al., 2009). For ChIP-qPCR, strains carried *REC8-3HA* (Klein et al., 1999), *SPO13-3FLAG*, *REC8-6HIS-3FLAG*, *SGO1-6HA*. To generate yeast strains carrying *SMC3* or *smc3K112R,K113R*, *SMC3* was amplified from genomic DNA and cloned into YIplac128 to generate AMp1342. A plasmid with *smc3-K112,113R* (AMp1392) was generated by site-directed mutagenesis (using QuikChange XL Site-directed mutagenesis kit, Agilent). Both plasmids were integrated into the *LEU2* locus in the *pCLB2-3HA-SMC3* strain background.

### Growth conditions

For asynchronous mitotic culture, cells were inoculated into YPDA (1% Bacto yeast extract, 2% Bacto peptone, 2% Glucose, 0.3 mM adenine) and grown at 30 °C with shaking at 250 rpm for ∼15 h. Cells were diluted to OD_600_=0.2 in YPDA and grown at room temperature for approximately 5 h until OD_600_ = 0.7-1.0. The OD_600_ was measured, and the cells were harvested for the experiment. For testing the viability of anchor-away strains YPDA agar plates contained 0.1 μM rapamycin.

For induction of meiosis, diploid yeasts were thawed onto YPG agar (1% Bacto yeast extract, 2% Bacto peptone, 2.5% glycerol, 0.3 mM adenine, 2% agar), and incubated at 30 °C overnight before transferring onto 4% YPDA agar (1% Bacto yeast extract, 2% Bacto peptone, 4% glucose, 0.3 mM adenine, 2% agar) and incubating at 30 °C 8h-overnight. Cells were inoculated into YPDA and shaken at 30 °C for 24 h, diluted into BYTA (1% Bacto yeast extract, 2% Bacto tryptone, 1% potassium acetate, 50 mM potassium phthalate) at OD_600_=0.2, and incubated at 30 °C with shaking at 250 rpm for 14-16 h. Cells were collected by centrifugation, washed twice in sterile dH_2_O, resuspended in SPO (0.3% potassium acetate, pH 7) at OD_600_ = 1.8-1.9 and incubated at 30 °C with shaking at 250 rpm to induce meiosis (t = 0). For synchronous induction of S phase, to cells carrying *pCUP1-IME1 pCUP1-IME4* (Berchowitz et al., 2013), 25 μM CuSO_4_ was added 2h after resuspension in SPO. For synchronous release from prophase, to cells carrying *pGAL1-NDT80 pGPD1-GAL4(848).ER* (Benjamin et al., 2003; Carlile and Amon, 2008) 1 μM β-estradiol was added 6h after resuspension in SPO.

For anchor-away experiments, all strains carried *tor1-1* and *fpr1Δ* mutations to prevent rapamycin toxicity in addition to Rpl13A-2xFKBP12. 1 μM rapamycin (5 mM rapamycin stock in DMSO (Dimethyl sulphoxide)) or DMSO (as a control) was added to all cultures at the time of resuspension in SPO.

### Spore viability

Diploid strains were sporulated on agar (>48h) or liquid (24h) SPO media (with addition of appropriate drugs) at 30 °C. Cells were treated with 20 μl 1 mg/ml zymolyase (AMS Biotechnology) in 2 M sorbitol for 8 min, before diluting with 1 ml dH_2_O. A minimum of 50 tetrads were dissected into individual spores on YPDA agar using a micromanipulator on a Nikon Eclipse 50i light microscope and viable colonies were scored ∼48h later.

### DNA staining

100 μl meiotic cell culture was added to 400 μl of 100 % EtOH and stored at 4 °C. For DNA staining, the cells were pelleted and resuspended in 20 μl of 1 μg/ml DAPI in PBS (13.7 mM NaCl, 270 μM KCl, 1 mM Na_2_PO_4_, 176 μM KH_2_PO_4_) and stored at 4 °C. For visualization and scoring, 3 μl of sample on a 1 mm thick glass slide (Fisher Scientific) and an 18 x18 mm cover slip (VWR) was imaged on a Zeiss Axioplan Imager Z2 fluorescence microscope with a 100 x Plan ApoChromat NA 1.45 oil lens. For each condition 200 cells were scored.

### Tubulin immunofluorescence

300 μl of meiotic cell culture was pelleted, resuspended in 500 μl of 3.7% formaldehyde solution (3.7% formaldehyde diluted in 0.1 M potassium phosphate buffer pH 6.4), and placed at 4 °C overnight. Cells were pelleted and washed three times by resuspending in 1 ml of 0.1 M potassium phosphate buffer pH 6.4, then pellets were resuspended in 1.2 M sorbitol-citrate (1.2 M sorbitol, 0.1 M K_2_HPO_4_, 36 mM citric acid). The cells were then either stored at -20 °C, or the immunofluorescence protocol immediately continued. Fixed, washed cells were pelleted, resuspended in 226 μl of Digestion Solution (200 μl 1.2 M sorbitol-citrate, 20 μl glusulase (PerkinElmer), 6 μl 10 mg/ml zymolyase (AMS Biotechnology)) and incubated at 30 °C for 2-3 h, until spheroplasts were observed by light microscopy. The digested cells were pelleted at 3000 rpm for 2 min, gently washed in 1.2 M sorbitol-citrate, resuspended in approximately 30 μl 1.2 M sorbitol-citrate before adhering 5 μl of cell suspension to each well of a multi-well slide (pre-treated by addition of 5 μl of 0.1% polylysine for 5 min, washing in dH_2_O and air-drying) for 10 min. The supernatant was aspirated, cell density confirmed by light microscopy, then the slide was incubated in 100% MeOH for 3 min followed by 10 sec in 100% acetone before air drying. To each well, 5 μl primary rat anti-tubulin antibody (Bio-Rad AbD Serotec) at a 1:50 dilution in PBS/1%BSA was added and the slide incubated in a dark wet chamber at room temperature for 2 h. Wells were washed individually five times in 5 μl PBS/BSA with aspiration between each wash. Secondary donkey anti-rat FITC antibody (Jackson ImmunoResearch) was added at a 1:100 dilution in PBS/BSA (5 μl), and incubated for a further 2 h before washing 5 times as above. DAPI-Mount (1 mg/ml *p*-phenylenediamine, 0.05 μg/ml DAPI, 40 mM K_2_HPO_4_, 10 mM KH_2_PO_4_, 150 mM NaCl, 0.1% NaN_3_, 90% glycerol) was added to each well (3 μl) and the slide was covered with a 24x60 mm cover-slip and sealed with nail varnish. Slides were then stored at - 20 °C before visualizing on a Zeiss Axioplan Imager Z2 fluorescence microscope with a 100 x Plan ApoChromat NA 1.45 oil lens. Spindle morphology was scored in 200 cells per timepoint.

### Flow cytometry

For flow cytometry, 500 μl of cycling or 150 μl of meiotic culture was fixed in 70% EtOH and stored at 4 °C. Pelleted cells were resuspended in 1 ml 50 mM Tris pH 7.5 and sonicated in a Bioruptor Twin (Diagenode) on LOW 30 sec ON. Pellets were collected, resuspended in 475 μl of 50 mM Tris-HCl pH 7.5 with 25 μl 20 mg/ml RNase A (Amresco) and incubated at 37 °C overnight. Cell pellets were washed in 1 ml 50 mM Tris pH 7.5, resuspended in 500 μl 50 mM Tris pH 7.5 with 10 μl 20 mg/ml Proteinase K (Invitrogen) and placed at 50 °C for 2 h. Collected cell pellets were washed in 1 ml 50 mM NaCitrate, resuspended in 500 μl 50 mM NaCitrate with 9.17 μl 1 mg/ml propridium iodide (Sigma Aldrich), and sonicated on LOW 30 sec ON, 30 sec OFF for 10 times. Samples were measured on a Becton Dickinson FACSCalibur with CellQuest Pro programme or an Attune NxT flow cytometer (n = 20,000 cells per sample) and analyzed using FlowJo V10. Signal intensity is shown on a linear scale.

### Visualization of GFP-labelled chromosomes

100 μl of culture was added to 10 μl of 37% formaldehyde and incubated for 8-9 min at room temperature before pelleting. The supernatant was aspirated, 1 ml of 80% EtOH added and tubes briefly vortexed. Cells were collected by short (∼15 sec) centrifugation, all traces of 80% EtOH removed by pipetting, the pellet was resuspended in 20 μl of 1 μg/ml DAPI in PBS and stored at 4 °C. For visualization, 3ul of sample was transferred to a glass slide, a coverslip added, pressure applied to flatten the cells before viewing under a Zeiss Axioplan Imager Z2 fluorescence microscope with a 100 x Plan ApoChromat NA 1.45 oil lens. All unbudded cells were scored for the presence of either one or two GFP dots (n=100). All experiments were done in triplicate.

### Live-cell imaging

8 -Well Glass Bottom µ-Slide Ibidi dishes (Ibidi) were prepared by spreading 45 μl 5 mg/ml concanavalin A (dissolved in 50 mM CaCl_2_, 50 mM MnCl_2_) evenly in the bottom of each well and incubating at 32 °C for 15 min. Excess concanavalin A was removed by aspiration, and each well washed three times in 500 μl dH_2_O. Around 2.30 h after meiotic cultures were resuspended in SPO (OD_600_=2.5) as described above, 3 ml of cell culture was harvested by centrifugation (3000 rpm, 3 min) to concentrate, resuspended in 300 μl of their supernatant (pre-conditioned), and added to the wells of the Ibidi dish. The dish was incubated at 30 °C for 20 min, the excess SPO culture aspirated and wells were washed twice in 500 μl dH_2_O and once in 500 μl of pre- conditioned SPO media, before adding 400 μl pre-conditioned SPO media to each well. Cells were visualized with a Zeiss Axio Observer Z1 (Zeiss UK, Cambridge) with a Prior motorized stage equipped with a Hamamatsu Flash 4 sCMOS camera and Zen 2.3 acquisition software on a Linux computer. Images were acquired every 15 min for 12 h, with 8 Z-stacks of 0.8 μM for FITC and Tomato channels and a single stack for Brightfield. For *CEN5*-GFP dots, FITC channel imaging conditions were: binning 2x2, 5 % transmitted light and 0.15 sec exposure. For imaging of Spc42-tdTomato and Pds1-tdTomato, the red channel imaging conditions were: binning 2x2, 5 % transmitted light and 0.2 sec exposure. The images were analyzed using the ImageJ software version 2.0.0-rc-43/1.51g (National Institutes of Health). For each experiment, all strains were imaged simultaneously in multi-well Ibidi dishes and experiments were performed at least twice, scoring at least 100 cells per strain. A representative experiment is shown.

### Western blotting

For mitotic protein extracts 10 ml of YPDA cell culture OD_600_=0.6-1 was collected, and for meiotic protein extracts 5 ml of SPO cell culture OD_600_=1.8 was collected. Cells were pelleted by centrifugation, resuspended in 5 ml 5% trichloroacetic acid and incubated on ice for a minimum of 10 min, before pelleting and transferring to a 2 ml fast-prep tube (MP Biomedicals). The fast-prep tube was centrifuged, the supernatant removed and snap-frozen in liquid nitrogen before storing at -80 °C. Thawed samples were resuspended in 1 ml acetone by vortexing, cells pelleted by centrifugation and the acetone removed. Samples were air-dried until the pellet was dry (>4h) and resuspended in 100 μl of protein breakage buffer (10 mM Tris-HCl pH 7.5, 1 mM EDTA pH 7.5, 2.75 mM DTT, 1 x Roche EDTA-free protease inhibitors) and one volume of glass beads (0.5mM zirconia/silica glass beads, Biospec Products) added. The cells were disrupted in a Fastprep Bio-Pulveriser FP120 at 6.5 speed for 3 cycles of 45 sec, placed on ice and 50 μl of 3xSDS sample buffer (187 mM Tris-HCl pH 6.8, 6% β-mercaptoethanol, 30% glycerol, 9% SDS, 0.05% bromophenol blue) was added to the lysate before immediately heating at 95 °C for 5 min, cooled and centrifuged before loading onto SDS-PAGE gels of the appropriate concentration (8-10%). PAGE was carried out using a Biorad Mini Trans-Blot System (Biorad) or a Biometra V17.15 System (Biometra) in SDS running buffer (25 mM Tris, 190 mM glycine, 0.01% SDS). SDS-PAGE gels were transferred onto nitrocellulose membrane (0.45 μM, GE Healthcare, Amersham) in transfer Buffer (25 mM Tris, 1.5% glycine, 0.02% SDS, 10% MeOH) in either a Biorad Mini Trans-Blot system or a semi-dry Amersham TE70 transfer unit. Membranes were blocked in 5 % milk in PBS with 0.05% Tween20 (PBST) for at least 1 h at room temperature before incubating in primary antibody in 2 % milk/PBST overnight at 4 °C. Membranes were washed in PBST three times for 15 min, incubated in secondary antibody in 2 % milk/PBST for 1 h at room temperature, washed in PBST three times. HRP-conjugated antibodies were detected with an ECL SuperSignal West Pico chemiluminescence kit (Thermo Scientific) ECL Supersignal West Pico, supplemented with SuperSignal West Femto chemiluminescence kit (Thermo Scientific) for weaker signals. Membranes were exposed to X-ray film (Agfa Healthcare CP-BU, blue), developed using a Konica-Minolta SRX-101A developer. Primary antibodies used were mouse anti-HA 12CA5 (Roche; 1:1000), mouse anti-HA 11 (Biolegend,; 1:1000), mouse anti-FLAG M2 (Sigma-; 1:1000), rabbit anti-Pgk1 (Marston lab stock; 1:10,000), rabbit anti-Kar2 (Marston lab stock; 1:10,000), rabbit anti Smc3-K112,113Ac (custom-made, Genescript; 1:1000), rabbit anti-Rec8 (Marston lab stock; 1:15,000), rabbit anti-Smc3 (Marston lab stock; 1:5000). Secondary antibodies used were sheep anti-mouse HRP (GE healthcare; 1:5000) and donkey anti-rabbit (GE healthcare; 1:5000).

### Chromatin Immunoprecipitation (ChIP) - qPCR

Meiotic culture (45 ml) was mixed with 5 ml of fixing solution (1.5 ml 37% formaldehyde in 3.5 ml diluent (143 mM NaCl, 1.43 mM EDTA, 71.43 mM Hepes-KOH pH 7.5)), and gently rocked at room temperature for 2h. Cells were pelleted and washed twice in 10 ml of ice-cold TBS (20 mM Tris-HCl pH 7.5, 150 mM NaCl), then once in 10 ml ice-cold 1xFA lysis buffer (50 mM Hepes-KOH pH 7.5, 150 mM NaCl, 1 mM EDTA, 1% Triton X-100, 0.1% Na Deoxycholate) with 0.1% SDS. Pellets were collected into 2 ml fast-prep tubes (MP Biomedicals), snap-frozen in liquid nitrogen and stored at -80 °C. Cell pellets were thawed on ice, and the pellet resuspended in 300 μl 1xFA lysis buffer with 0.5% SDS, 1x Roche EDTA-free protease inhibitors, 1 mM PMSF and one scoop of glass beads (0.5mM zirconia/silica glass beads, Biospec Products) was added. Cells were disrupted in a Fastprep Bio-Pulveriser FP120 at 6.5 speed for 2 cycles of 30 sec, with a 10 min waiting period on ice between. The tube was punctured, the cell lysate and debris were collected by centrifugation, transferred to a new tube and then pelleted by centrifugation at 13,000 rpm for 15 min at 4 °C. The cell pellet was resuspended in 1 ml 1xFA lysis buffer with 0.1% SDS, 1x Roche EDTA-free protease inhibitors and 1 mM PMSF, and re-centrifuged. The supernatant was removed, the pellet resuspended in 300 μl 1xFA lysis buffer with 0.1% SDS, 1x Roche EDTA-free protease inhibitors and 1 mM PMSF, then sonicated at 4 °C in a Bioruptor Twin sonicating device (Diagenode) on HIGH 30 sec ON, 30 sec OFF for 30 min total. The cell debris was pelleted at 13,000 rpm for 15 min at 4 °C, and the supernatant transferred into a new 1.5 ml tube containing 1 ml of 1xFA lysis buffer with 0.1% SDS, 1x Roche EDTA-free protease inhibitors and 1 mM PMSF. The chromatin lysate was re-centrifuged, and the supernatant transferred into a new tube. For the INPUT sample, 10 μl of the supernatant was frozen at -20 °C overnight. For the IP, 15 μl of 20 mg/ml Protein G Dynabeads (Invitrogen) per sample were pre-washed four times in 1 ml 1xFA lysis buffer with 0.1% SDS, 1x Roche EDTA-free protease inhibitors and 1 mM PMSF, then resuspended in 100ul/sample 1xFA lysis buffer with 0.1% SDS, 1x Roche EDTA-free protease inhibitors and 1 mM PMSF. To 1 ml of cell supernatant, 100 μl of Protein G Dynabeads was added, along with the appropriate antibody (mouse HA 12CA5, 7.5 μl 0.4 mg/ml, Roche; mouse M2 FLAG, 5 μl 1 mg/ml, Sigma), before incubating at 4 °C on a rotating wheel at 14 rpm for 15-18 h. Following incubation, the beads were collected on a magnet, and the supernatant discarded. The beads were washed 5 min/wash consecutively in ChIP wash buffer 1 (1xFA lysis buffer, 0.1% SDS, 275 mM NaCl), ChIP wash buffer 2 (1xFA lysis buffer, 0.1% SDS, 500 mM NaCl), ChIP wash buffer 3 (10 mM Tris-HCl pH 8, 250 mM LiCl, 1 mM EDTA, 0.5% NP-40, 0.5% Na Deoxycholate), and ChIP wash buffer 4 (10 mM Tris-HCl pH 8, 1 mM EDTA), with 30 sec on a small magnet in-between each wash. After the final wash, all of the supernatant was removed from the beads.

Chelex-100 Resin (Biorad) was resuspended at 0.1 g/ml in Hyclone water (Hypure Molecular Biology Grade Water, GE Lifesciences), and 100 μl added to both the thawed and vortexed INPUT samples and the IP samples. The samples were boiled at 100 °C for 10 min, cooled on ice, then briefly centrifuged at 2000 rpm for 1 min. To each tube 2.5 μl 10 mg/ml Proteinase K (Invitrogen) was added, the samples vortexed and then incubated at 55 °C for 30 min. The samples were again boiled at 100 °C for 10 min, cooled on ice, then briefly centrifuged at 2000 rpm for 1 min. Approximately 120 μl of supernatant was transferred into a new tube, and frozen at -20 °C. For qPCR either SYBR GreenER (Invitrogen; Figure S5B-C) or Luna (NEB; Figure S5E) mastermix was used. Input DNA was diluted 1:500 or 1:300 and the ChIP DNA was diluted 1:10 or 1: 6 in Hyclone Water for SYBR GreenER or Luna, respectively. For each biological replicate, qPCR reactions were carried out in technical triplicate on a Lightcycler 480 Roche machine with 40 cycles for SYBR GreenER and 45 cycles for NEB Luna. The threshold cycle (Ct) values were computed by the Lightcycler 480 Roche software using the 2^nd^ derivative maximum algorithm. The geometric mean of technical replicate Ct values was determined and delta Ct was calculated according to the following formula: ΔCt = Ct_(ChIP)_ (Ct_(Input)_ log_(primer efficiency)_(Input dilution factor)). Enrichment values are given as ChIP/Input = (primer efficiency)^^(-ΔCt)^. Primers used are given in Table S3.

### ChIP - sequencing

For calibrated ChIP-seq (Smc3) we used the method of (Hu et al., 2015), as modified by (Galander et al., 2019b). For each condition 200 ml of *S. cerevisiae* meiotic culture was grown and fixed as for ChIP-qPCR and frozen in four pellets (each from 50 ml culture). Each pellet was mixed with a fixed pellet from 100 ml *S. pombe* culture prepared as follows. Wild type *S. pombe* (strain spAM29) was inoculated into YES media (0.5% yeast extract, 3% glucose, 2% agar 225mg/l each of adenine, histidine, leucin, uracil and lysine hydrochloride), grown for 16 h at 30 °C before diluting to OD_600_=0.07-0.1 and growing for approximately 4-6 h at 30 °C until the OD_600_=0.4. Cell culture (1l) was mixed with 100 ml of fixing solution (33.3 ml 37% formaldehyde in 66.6 ml diluent) and incubated at room temperature at 90 rpm for 2h. Cells were harvested, washed twice with TBS and completely resuspended in 1xFA lysis/0.1% SDS. An equal amount of cells, equivalent to 100 ml of culture, was evenly distributed into Fastprep tubes and snap frozen. Thawed *S. cerevisiae* and *S. pombe* cell pellets were combined on ice in 400 μl 1xFA lysis buffer with 0.5% SDS, 1x Roche EDTA-free protease inhibitors and 1 mM PMSF. Cells were processed as for ChIP-qPCR except 4 cycles of 60 sec on Fastprep Bio-Pulveriser FP120 and two rounds of 30 cycles 30 sec ON-30sec OFF of sonication with Bioruptor Plus (Diagenode) set on HIGH were performed. Before setting up the IP, the four samples of each condition were pooled together and then split into a 10 μl INPUT and 4 x 1ml IPs, to whom 15 μl of Protein G Dynabeads (Invitrogen) and 10 μl of Rabbit anti-Smc3 (Marston lab stock) were added. After overnight incubation, washes were performed as for ChIP-qPCR and after the final wash the beads for each condition were pooled in 200 μl TES (50 mM Tris-HCl pH 7.5, 10 mM EDTA, 1% SDS), the sample eluted at 65 °C for 20 min and transferred to a new tube. The beads were washed for 15 min at room temperature in 200 μl TE (10 mM Tris-HCl pH 7.5, 1 mM EDTA pH 7.5), that was then combined with the corresponding eluate. 40 μl 10 mg/ml Proteinase K was added, and the samples were incubated at 42 °C for 1 h then at 65 °C for 16 h. Samples were cleaned up using a Promega Wizard Kit, eluted in 35 μl Hyclone water and frozen at -20 °C.

Non-calibrated ChIP-seq (Rec8) samples were processed with the same steps and conditions as the calibrated ones, but without adding *S. pombe* cell pellets.

ChIP-sequencing libraries were prepared using NEXTflex-6 DNA Barcodes (PerkinElmer) in DNA LoBind tubes (Eppendorf) using Hyclone water. Briefly, 2 ng of INPUT or IP DNA was taken, and blunt and phosphorylated ends were generated using the Quick Blunting kit (NEB), before removal of DNA fragments under 100 bp using AMPure XP beads (Beckman Coulter), followed by addition of dA tails by Klenow fragment (*exo-*) enzyme (NEB). Suitable NEXTflex-6 barcodes were ligated using Quick Ligation kit (NEB) before consecutive selection of DNA fragments >100 bp and then >150-200 bp using AMPure XP beads. Libraries were amplified by PCR using NextFlex PCR primers (Primer 1 - 5’-AATGATACGGCGACCACCGAGATCTACAC; Primer 2 - 5’-CAAGCAGAAGACGGCATACGAGAT) and Phusion High-Fidelity DNA Polymerase (NEB) before three further rounds of AMPure XP purification were performed to collect fragments 150-300 bp in size. Library quality was assessed on a Bioanalyzer (Agilent) using the 2100 Bioanalyzer High Sensitivity DNA kit (Agilent) and DNA was quantified by QuBit (Thermo Scientific), before preparation and denaturation of a 1 nM pooled library sample where INPUTs and IPs were mixed in 15%-85% ratio, followed by sequencing in house on an Illumina MiniSeq instrument (Illumina) with Miniseq High output reagent kit (150-cycles) (Illumina). Calculation of Occupancy Ratio (OR) and data analysis was performed as described by (Hu et al., 2015). Briefly, reads were mapped to both the *S. pombe* calibration and *S. cerevisiae* SK1 experimental genomes and the number of reads mapping to each genome was determined. Occupancy ratio was then determined using the formula Wc*IPx/Wx*IPc where W=Input; IP=ChIP; c=calibration genome (*S. pombe*) and x=experimental genome (*S. cerevisiae* SK1). The number of reads at each position was normalized using the OR and visualized using the Integrated Genome Viewer (IGV, Broad Institute). Mean calibrated ChIP-Seq read plots (pile-up) at centromeres, pericentromeric borders and arm sites were generated using the Bioconductor SeqPlots package. Reads were binned at 50bp windows around midpoint of centromeres/pericentromeric borders/arms with 3kb flanks at either side. Pericentromeric borders were oriented so that their position relative to the centromere was the same. To obtain genomic coordinates of borders in SK1, border cohesin peaks corresponding to those defined in w303 (Paldi et al., 2020) were identified from wild type Smc3 ChIP-Seq tracts visualized in IGV (either homologous sequence or, where the sequence was divergent, the position between the same gene pair). For the plots of cohesin enrichment on chromosome arms, the coordinates were chosen as for pericentromere borders; when W303 coordinates did not correspond to a cohesin peak in SK1, the next peak (distal from the centromere) was chosen. Centromere, pericentromere border and arm peak coordinates are listed in Table S4.

### Hi-C

The Hi-C protocol was adapted from (Paldi et al., 2020; Schalbetter et al., 2019). Briefly, 50 ml of synchronized meiotic culture at OD_600_∼2 was fixed for 20 min in 3% formaldehyde at room temperature with 90 rpm shaking. The reaction was quenched by adding Glycine to 0.35 M and incubating for 5 min. Cells were harvested and washed once in 40 ml of cold water, resuspended in 5 ml 1x NEB Buffer 2 (NEB) and drop frozen in liquid nitrogen. Frozen pellets were ground in a chilled pestle and mortar on dry ice for 20 min with addition of liquid nitrogen every 3 min. About 0.5g of crushed pellets (adjusted to the measured culture OD_600_) were then thawed, washed with 1x NEB Buffer 3.1 (NEB) and digested with 2 U/μl of DpnII restriction nuclease (NEB) at 37 °C overnight. Digested DNA ends were filled-in and biotinylated with a nucleotide mix containing 0.4 mM biotin-14-dCTP (Invitrogen) instead of dCTP and 5 U/μl Klenow fragment Dpol I (NEB) at 37°C for 2 h. The reaction was stopped by incubating at 65 °C for 20 min with 1.5% SDS. Proximity ligation was then carried out on diluted samples (25 ml) with 0.024 U/μl of T4 DNA ligase (Invitrogen) at 16 °C for 8 h, after which the ligase was inactivated with 5.3 mM EDTA pH8.0. DNA was decrosslinked overnight at 65°C with 70 ug/μl proteinase K (Invitrogen), topped up to 140 ug/ul for the final 2 h and then DNA was extracted with 20 ml Phenol:Chloroform:Isoamyl Alcohol 25:24:1 (Sigma), precipitated with ethanol, and resuspended in 22.5 ml TE buffer pH8.0. To concentrate and desalt DNA, samples were filtered through Amicon 30 kDa columns (Merck). DNA was then subjected to a second round of phenol extraction and ethanol purification, before being resuspended in 20 μl of TE and treated first with 1 mg/ml RNAse A (Amresco) for 1 h at 37 °C, next with 3U of T4 DNA Polymerase (NEB) for each ug of DNA and 5 mM dNTPs for 4 h at 20 °C to remove biotin from unligated ends. DNA was fragmented with 2 rounds of 30 cycles 30sec ON-30sec OFF of sonication with Bioruptor Plus (Diagenode) set on HIGH and purified with Qiagen MinElute kit (Qiagen). After fragmentation, DNA ends were repaired by treatment with 3U T4 DNA polymerase (NEB), 50U T4 Polynucleotide Kinase (NEB) and 25U Klenow fragment DPol I (NEB) for 30 min at 20 °C. DNA was once again purified with a Qiagen MinElute kit and dA-tailed with 11.25U Klenow fragment (*exo-*) enzyme (NEB). DNA fragments were then selected for size (200-300bp) with two subsequent rounds of AMPure XP bead (Beckman) treatment and for the presence of the Biotin marker by Dynabeads MyOne Streptavidin C1 beads (Invitrogen) pull-down. Finally, NEXTflex-6 barcoded adapters (PerkinElmer) were ligated using 1 μl Quick T4 DNA Ligase (NEB) and DNA fragments were amplified by PCR with NEXTFlex primers (Primer 1 - 5’-AATGATACGGCGACCACCGAGATCTACAC; Primer 2 - 5’-CAAGCAGAAGACGGCATACGAGAT) and Phusion High-Fidelity DNA Polymerase (NEB) and further purified with AMPure XP beads to eliminate unligated primers and adapters. Libraries were sequenced with 42 bp paired-end reads on an Illumina NextSeq500 (EMBL Core Genomics Facility, Heidelberg, Germany). Hi-C read number for each library are listed in Table S5.

For Hi-C data analysis, Fastq reads were aligned to *S. cerevisiae* SK1 reference genome using HiC-Pro v2.11.4 bowtie2 v2.3.4.1 (--very-sensitive -L 30 --score-min L,-0.6,-0.2 --end-to-end --reorder), removing singleton, multi hit, duplicated and MAPQ<10 reads. Read pairs were assigned to restriction fragment (*Dpn*II) and invalid pairs filtered out. Valid interaction pairs were converted into the .cool contact matrix format using the cooler library, and matrixes balanced using Iterative correction down to one kilobase resolution. Multi-resolution cool files were uploaded onto a local HiGlass server. To generate pile-ups at centromeres/pericentromeric borders, the cooltools library was used and cool matrices were binned at one kilobase resolutions. Plots were created around the midpoint of centromeres/pericentromeric borders, at the same coordinates used for ChIP-seq plots (listed in Table S4), with 25, 50, 100 kb flanks on each side, showing the log10 mean interaction frequency using a colour map similar to HiGlass ‘fall’. All centromere annotations were duplicated in both the forward/reverse strand orientations to create an average image which is mirror symmetrical. Pile-ups at pericentromeric borders were oriented where positions are identical in relation to the centromere. The ratio pile-ups between samples were created in a similar fashion plotting the log2 difference between samples in the ‘coolwarm’ colour map, e.g. A/B; red signifying increased contacts in A relative to B and blue decreased contacts in B relative to A. Scripts for visualisation will be available at (https://github.com/danrobertson87/Barton_2021). Cooler ‘show’ was also used to generate individual plots for each chromosome. Contact probability P(s) and its derivative slope plots were generated using the cooltools library. White stripes on plots represent regions where data was lost stochastically during mapping due to stringency settings filtering out reads. Hi-C matrices were aligned to ChIP-seq tracks on HiGlass.

