## Supplementary information for "Eco1-dependent cohesin acetylation anchors chromatin loops and cohesion to define functional meiotic chromosome domains"

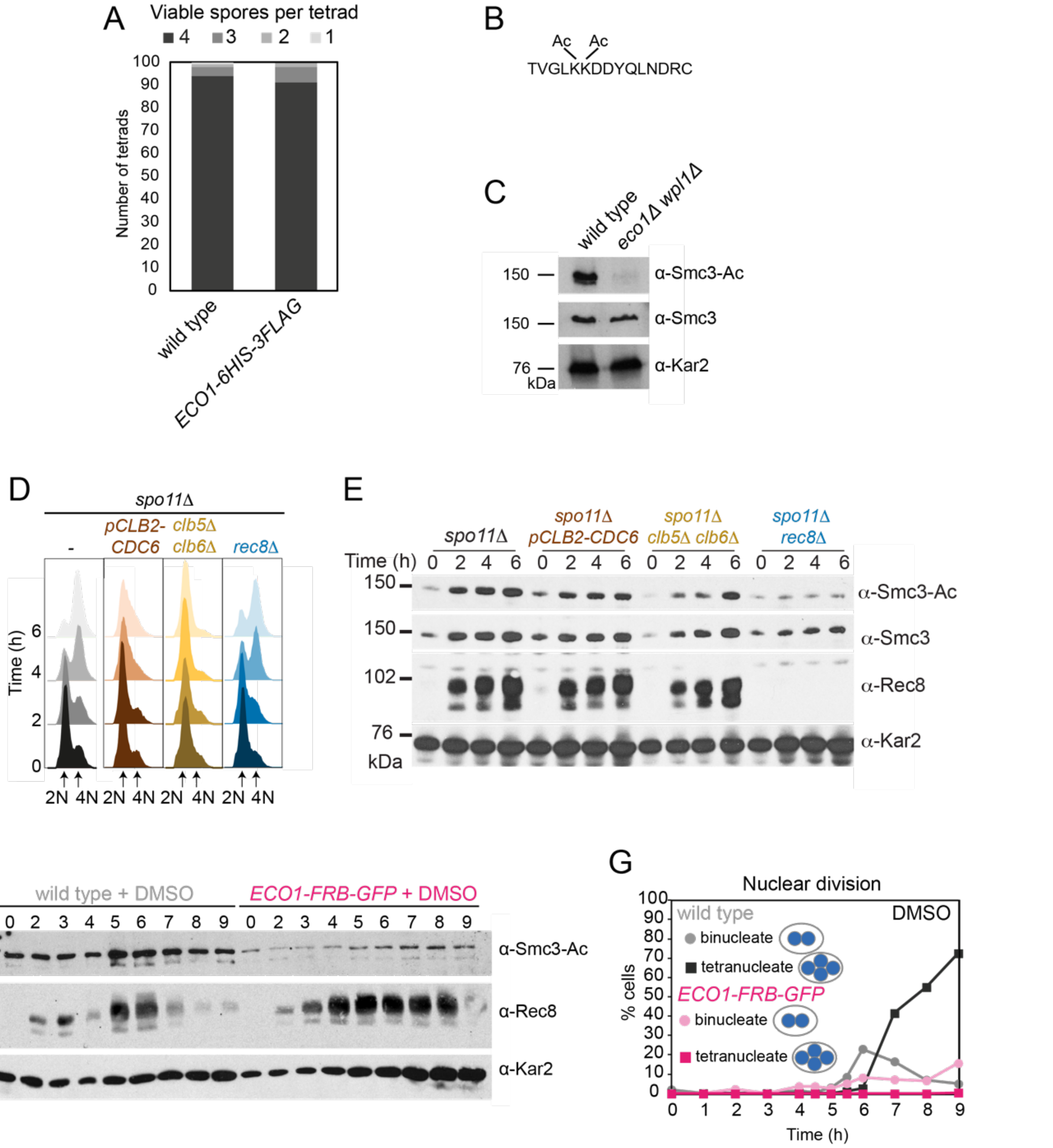


**Supplemental Figure S1. Tools to analyze Eco1-dependent Smc3 acetylation in meiosis**
(A) Eco1-6HIS-3FLAG is functional in meiosis. Spore viability of wild type (AM1835) and *ECO1-6HIS-3FLAG* (AM25844) strains. (B and C) An antibody to specifically recognise acetylated Smc3-K112,K113 in budding yeast. (B) Sequence of the peptide used as an immunogen for α-Smc3-Ac antibody generation. (C) Western immunoblot showing specificity of α-Smc3-Ac. Exponentially growing wild type (AM1176) and *eco1Δ wpl1Δ* (AM5433) cells were collected and total protein extracts analyzed by immunoblotting with α-Smc3-Ac, α-Smc3 and α-Kar2 antibodies. (D and E) Smc3-K112,113 acetylation on unreplicated chromosomes is not a consequence of double strand break formation during meiotic recombination or meiotic replication. *spo11Δ* (AM28949), *spo11Δ pSCC1-CDC6* (AM28948), *spo11Δ clb5Δ clb6Δ* (AM28947) and *spo11Δ rec8Δ* (AM28950) cells carrying *ndt80Δ* were analyzed as in Figure 1D and E. Flow cytometry profiles (D) and western immunoblots (E) against Smc3-Ac (α-Smc3-K112,113-Ac), Smc3 (α-Smc3), Rec8 (α-Rec8) and Kar2 loading control (α-Kar2) are shown. (F and G) Eco1-FRB-GFP is non-functional in meiosis even in the absence of rapamycin**.** Anchor away wild type (AM25532) and *eco1-aa* (AM22034) cells were induced to sporulate as in Figure 1 (H and I), except that DMSO was added rather than rapamycin. (F) Western immunoblot of whole cell extracts showing Smc3-Ac (α-Smc3-K112,113-Ac), Rec8 (α-Rec8), and Kar2 loading control (α-Kar2). (G) The percentages of bi- and tetra-nucleate cells were scored after DAPI-staining at the indicated timepoints (n = 200 cells/time point).

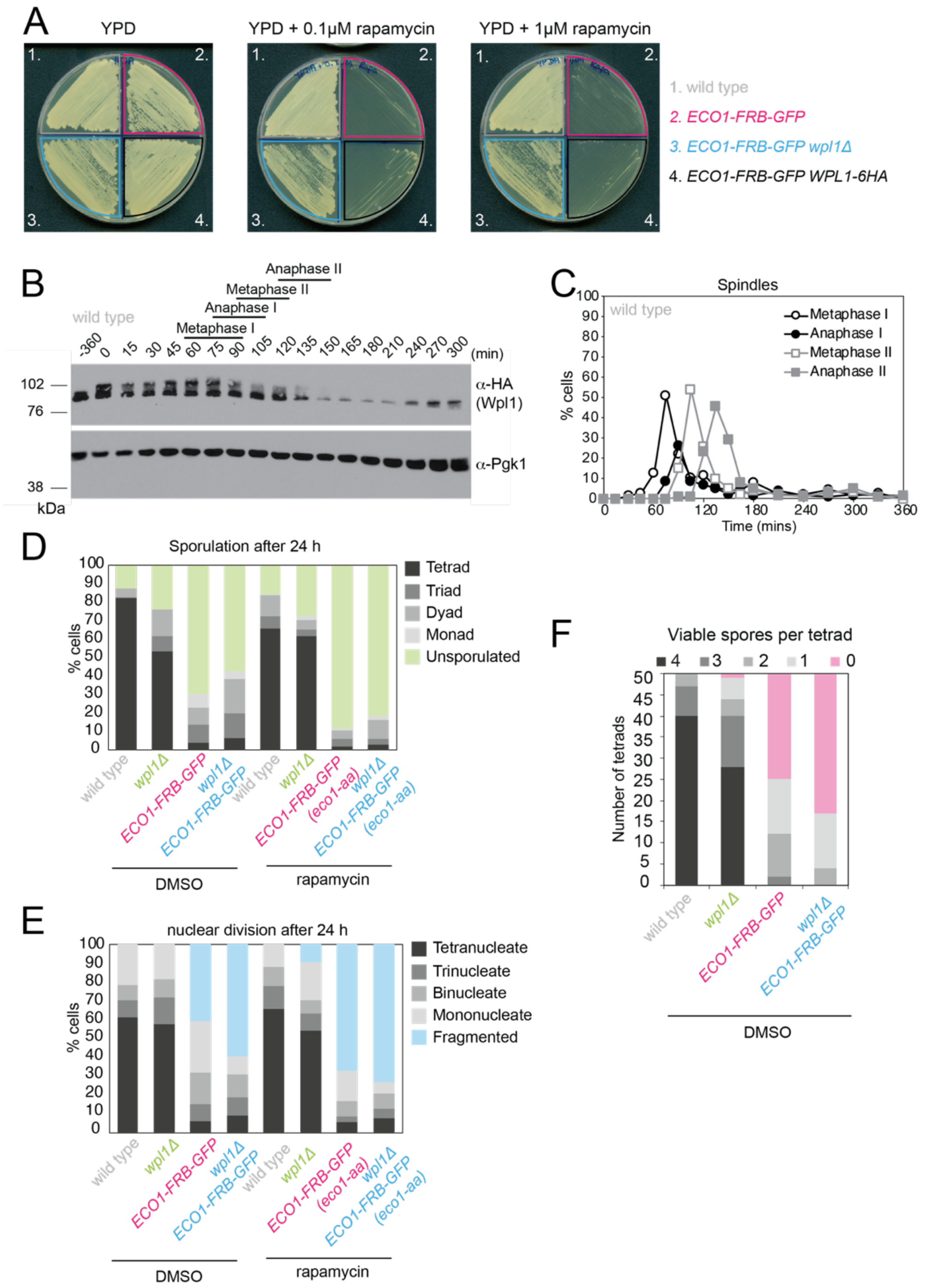


**Supplemental Figure S2. Wpl1-6HA is functional and Eco1 is required for meiosis even in the absence of Wpl1.**

(A) Diploid anchor-away wild type (AM23130), *eco1-aa* (AM22998), *eco1-aa wpl1Δ* (AM23415) and *eco1-aa WPL1-6HA* (AM23161) strains were patched onto either YPD, YPD + 0.1 μM rapamycin, or YPD + 1 μM rapamycin agar plates and incubated at 30ºC for 48 h. (B and C) Cells carrying *WPL1-6HA* and the *pGAL-NDT80* block/release genetic background (strain AM20953) were induced to sporulate. After 6 h, 1 μM β-estradiol was added to induce *NDT80* expression and release cells from prophase arrest. (B) Western immunoblot showing Wpl1-6HA (α-HA) and Pgk1 loading control (α-Pgk1). (C) Meiotic progression was monitored by scoring spindle morphology after α-tubulin immunofluorescence (*n=200* per timepoint). (D-F) Wild type, *eco1-aa*, *wpl1Δ* and *eco1-aa wpl1Δ* cells as in Figure 2D were sporulated in the presence of rapamycin or DMSO. (D) Sporulation efficiency was determined after 24h (*n=200*). (E) The number and morphology of nuclei was scored (*n=200*). (F) Spore viability was determined as described in Figure 2D, except that cells were sporulated in the presence of DMSO rather than rapamycin.

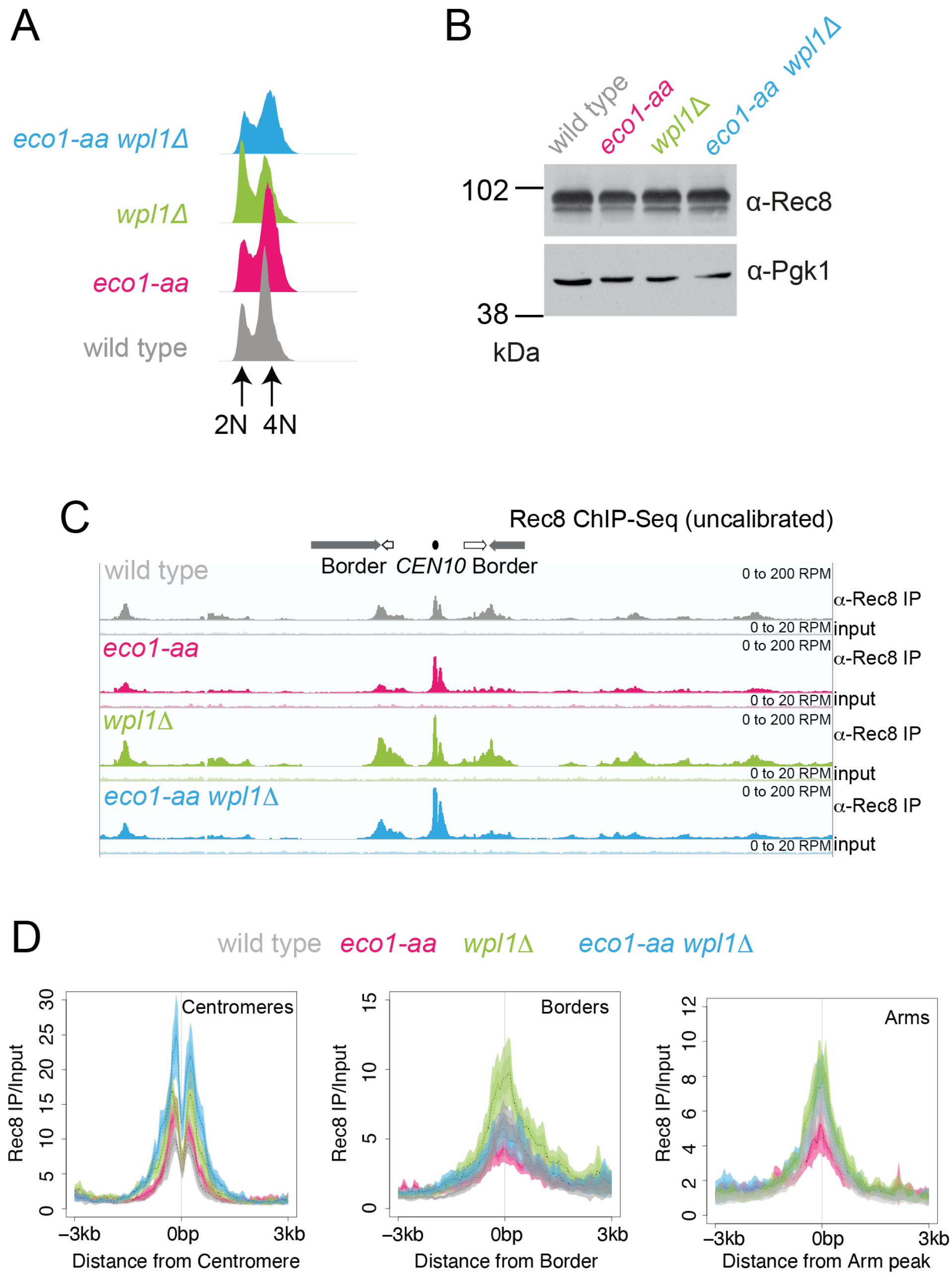


**Supplemental Figure S3. Rec8 levels at pericentromere borders, but not centromeres, require an Eco1 activity independent of Wpl1 antagonism.**
Rec8 ChIP-Seq was performed in prophase I using the strains as described in Figure 3. (A) Flow cytometry profiles show similar DNA content at harvesting in all cultures (B) Western immunoblot with Pgk1 loading control (α-Pgk1) shows comparable Rec8 (α-Rec8) levels in all cultures at the time of harvesting. (C) Calibrated Rec8 ChIP-seq for a representative region surrounding *CEN10*. (D) Mean calibrated ChIP-Seq reads (line), standard error (dark shading) and 95% confidence interval (light shading) at all 16 centromeres, 32 borders or 32 arm sites.

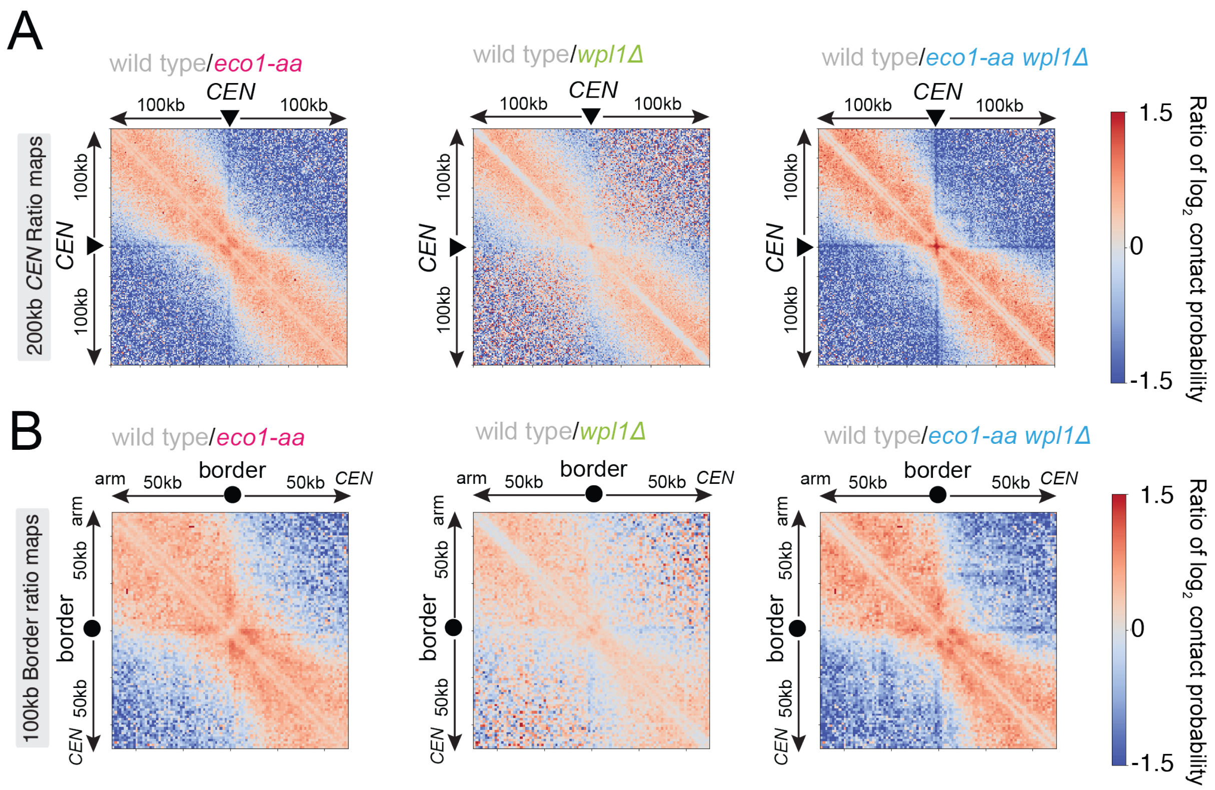


**Supplemental Figure S4. Eco1 is required for loop strength and positioning in meiotic prophase.**
Ratio plots of $log_2$ contact probability *CEN* and border pile-ups for the Hi-C data shown in Figure 4. Pair-wise comparisons of each of *eco1-aa*, *wpl1Δ* and *eco1-aa wpl1Δ* with the wild type are shown for the (A) 200kb region flanking the centromeres and (B) 100kb region flanking the borders.

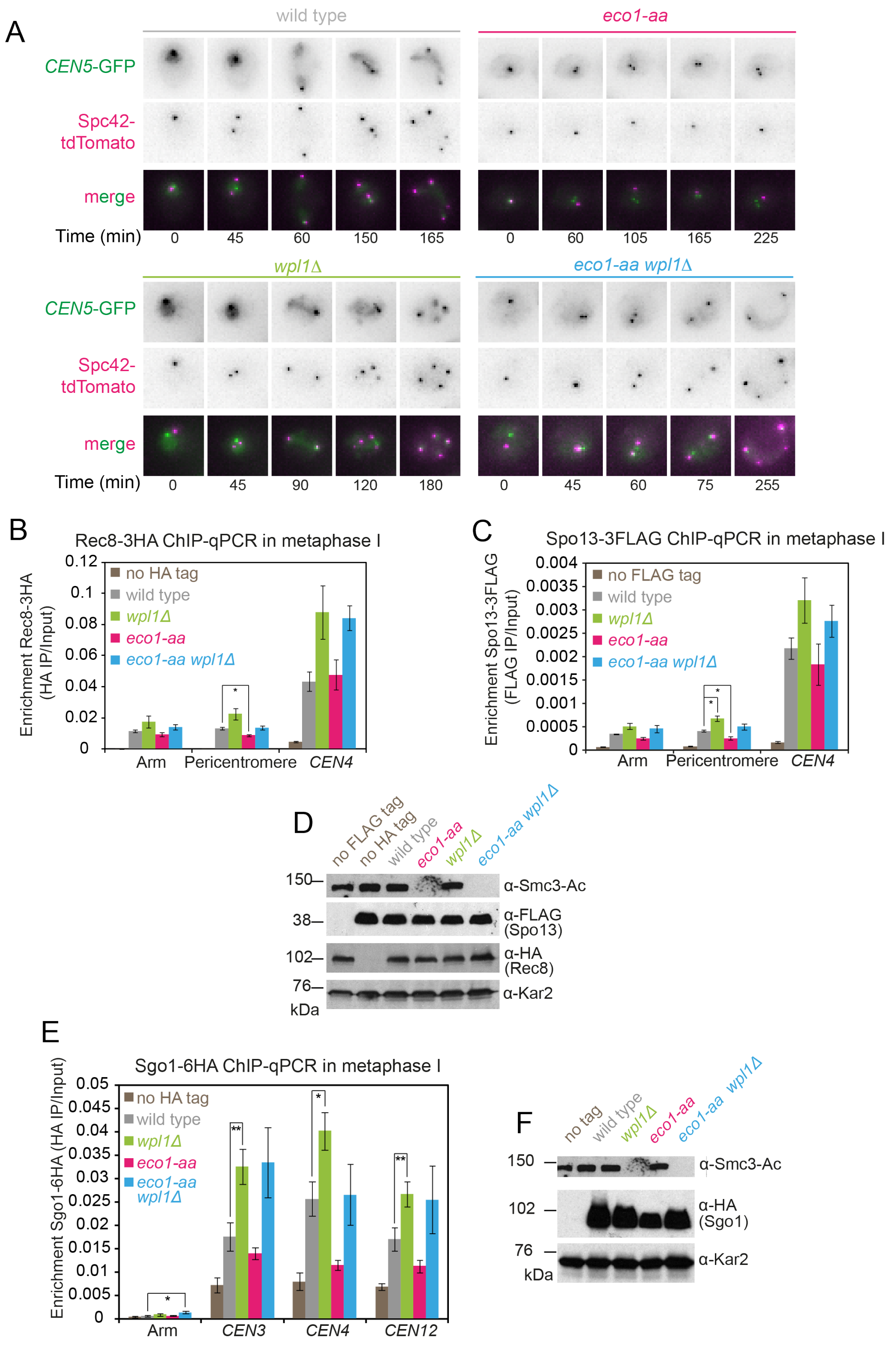


**Supplemental Figure S5. Sister kinetochore monoorientation and pericentromeric cohesion defects in *eco1-aa* cells are not a consequence of *SPO11* deletion or cohesin protector proteins Spo13 or Sgo1 mis-localization.**

(A) Representative images of wild type (AM24167), *wpl1Δ* (AM24168), *eco1-aa* (AM24184) and *eco1-aa wpl1Δ* (AM24169) anchor away cells carrying heterozygous *CEN5*-GFP and *SPC42*-tdTomato. Cells were imaged in the presence of rapamycin together with *spo11Δ* background strains in the experiment shown in Figure 6B-E. (B-D) Cohesin protector Spo13 localization follows the profile of Rec8 in metaphase I *eco1-aa* and *wpl1Δ* single and double mutant cells. Diploid strains *wpl1Δ* (AM24263), *eco1-aa* (AM24262) and *wpl1Δ eco1-aa* (AM24261) carrying *RPL13A-FKBP12, fpr1Δ, tor1-1, pCLB2-CDC20, REC8-3HA* and *SPO13-3FLAG* and wild type anchor away diploid strains carrying either *REC8-3HA* (AM24236) or *SPO13-3FLAG* (AM24235) were induced to sporulate in 1 μM rapamycin and harvested after 6 h. (B and C) Chromatin-bound Rec8 and Spo13 are increased in *wpl1Δ* and *eco1-aa wpl1Δ* mutants. Mean α-HA (B; Rec8-3HA) or α-FLAG (C, Spo13-3FLAG) ChIP-qPCR for the indicated sites is shown from 4 repeats with error bars representing standard error. * p <0.05, paired student T test. (D) Western immunoblot using α-Smc3-K112,113-Ac, α-FLAG, α-HA and α-Kar2 (loading control) confirming similar total levels of Rec8 and Spo13 in the expected strains. (E and F) Reduction in pericentromeric Sgo1 cannot explain the segregation defect of *wpl1Δ eco1-aa* cells. Wild type (AM24054), *wpl1Δ* (AM25103), *eco1-aa* (AM25104), and *eco1-aa wpl1Δ* (AM25102) strains carrying *RPL13A-FKBP12, fpr1Δ, tor1-1, pCLB2-CDC20, SGO1-6HA*, as well as a no tag strain (AM23700) without *SGO1-6HA*, were induced to sporulate in 1 μM rapamycin and harvested after 6 h (metaphase I). (E) Mean α-HA ChIP-qPCR (Sgo1-6HA) is shown for 4 replicates at the indicated sites with error bars showing standard error. **p<0.01, *p<0.05, paired student t test. (F) Western immunoblot probed with α-Smc3-K112,113-Ac, α-HA (Sgo1-6HA) and α-Kar2 (loading control).

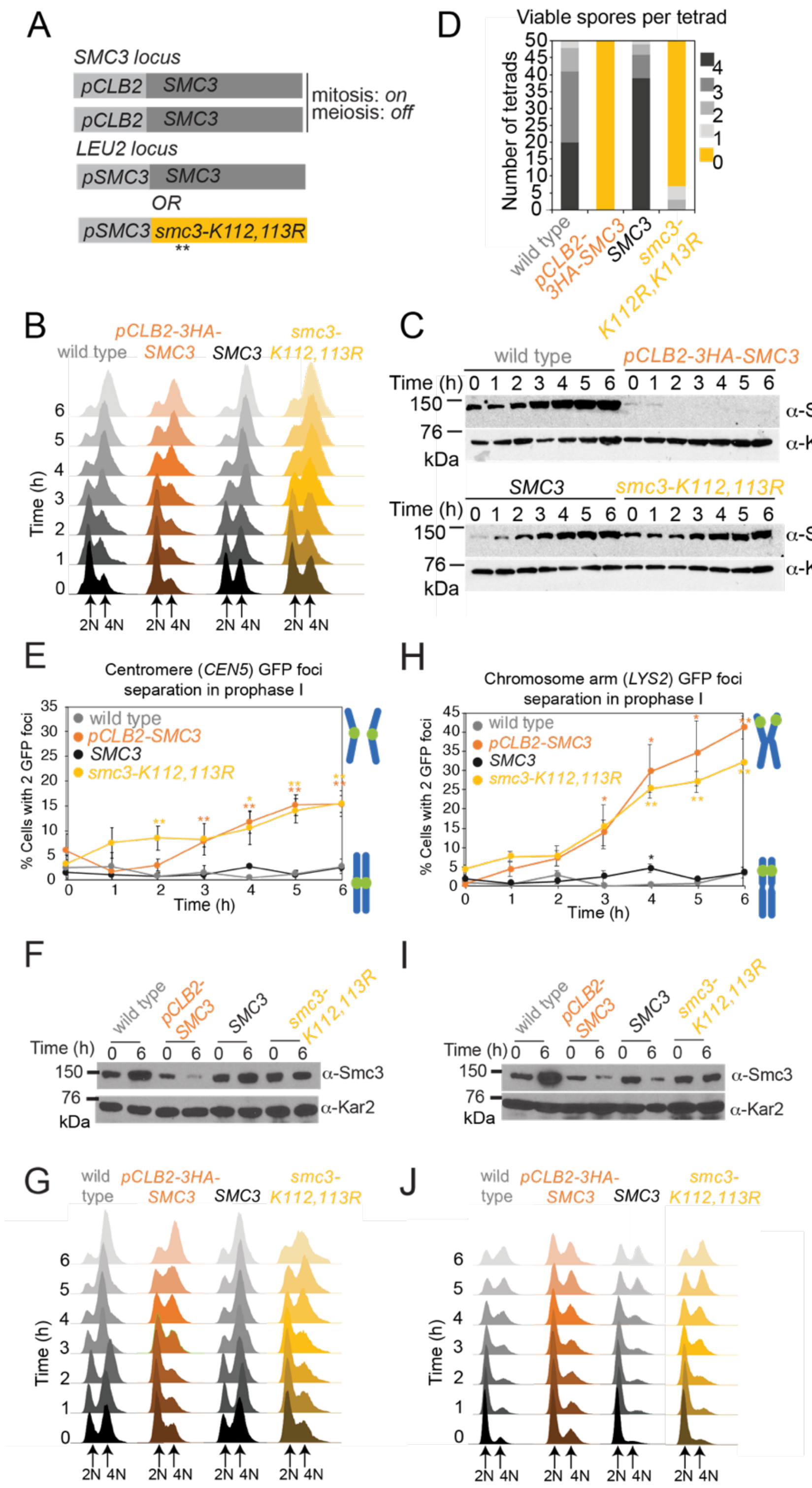


**Supplemental Figure S6. Chromosome segregation defects in *smc3-K112,113R* cells are modest, likely due to residual mitotic Smc3 expressed from *pCLB2-SMC3***

(A-C) Ectopic expression of *SMC3* and *SMC3-K112,113R* in meiosis. (A) Strategy to express non-acetylatable Smc3-K112,113R as the only Smc3 source in meiosis. In all diploid strains, both copies of endogenous *SMC3* are placed under control of the *CLB2* promoter, which is repressed in meiosis. Either wild type *SMC3* or *smc3-K112,113R* under the endogenous promoter is integrated at an ectopic locus of one parent (heterozygous). (B and C) Smc3-K112,113R is expressed but residual Smc3 persists from the *pCLB2-SMC3* construct after meiotic induction. Wild type (AM11633), *pCLB2-SMC3* (AM28718), *SMC3* (AM29315) and *smc3-K112,113R* (AM29316) carrying *ndt80Δ* were induced to sporulate. (B) Flow cytometry confirms DNA replication and prophase arrest. (C) Western immunoblot shows Smc3 ($\alpha$-Smc3) and Kar2 ($\alpha$-Kar2, loading control) levels at the indicated times after inducing sporulation. (D) Smc3 acetylation is required for meiosis. Spore viability of wild type (AM25765), *pCLB2-3HA-SMC3* (AM25067), *SMC3* (AM25920) and *smc3-K112,113R* (AM28741) cells. (E-J) Establishment of meiotic cohesion requires Smc3 acetylation. Cells carrying *ndt80Δ* and heterozygous *CEN5*-GFP (E-G) or *LYS2*-GFP (H-J) were induced to sporulate. The mean percentage of cells with two visible *CEN5*-GFP (E) or *LYS2*-GFP (H) foci were scored at the indicated timepoints in four (E) or three (H) biological replicates (100 cells per timepoint). Bars show standard error. * $p<0.05$; ** $p<0.005$, paired t test. Western immunoblots (F and I) show Smc3 ($\alpha$-Smc3) and loading control ($\alpha$-Kar2) levels at the indicated timepoints for a representative experiment. (G and J) Flow cytometry shows DNA content at the indicated timepoints for a representative experiment. Strains used in (E-G) were wild type (AM29184), *pCLB2-3HA-SMC3* (AM29185), *SMC3* (AM29186) or *smc3-K112,113R* (AM29187). Strains used in (H-J) were wild type (AM29247), *pCLB2-3HA-SMC3* (AM29248), *SMC3* (AM29249) or *smc3-K112,113R* (AM29250).

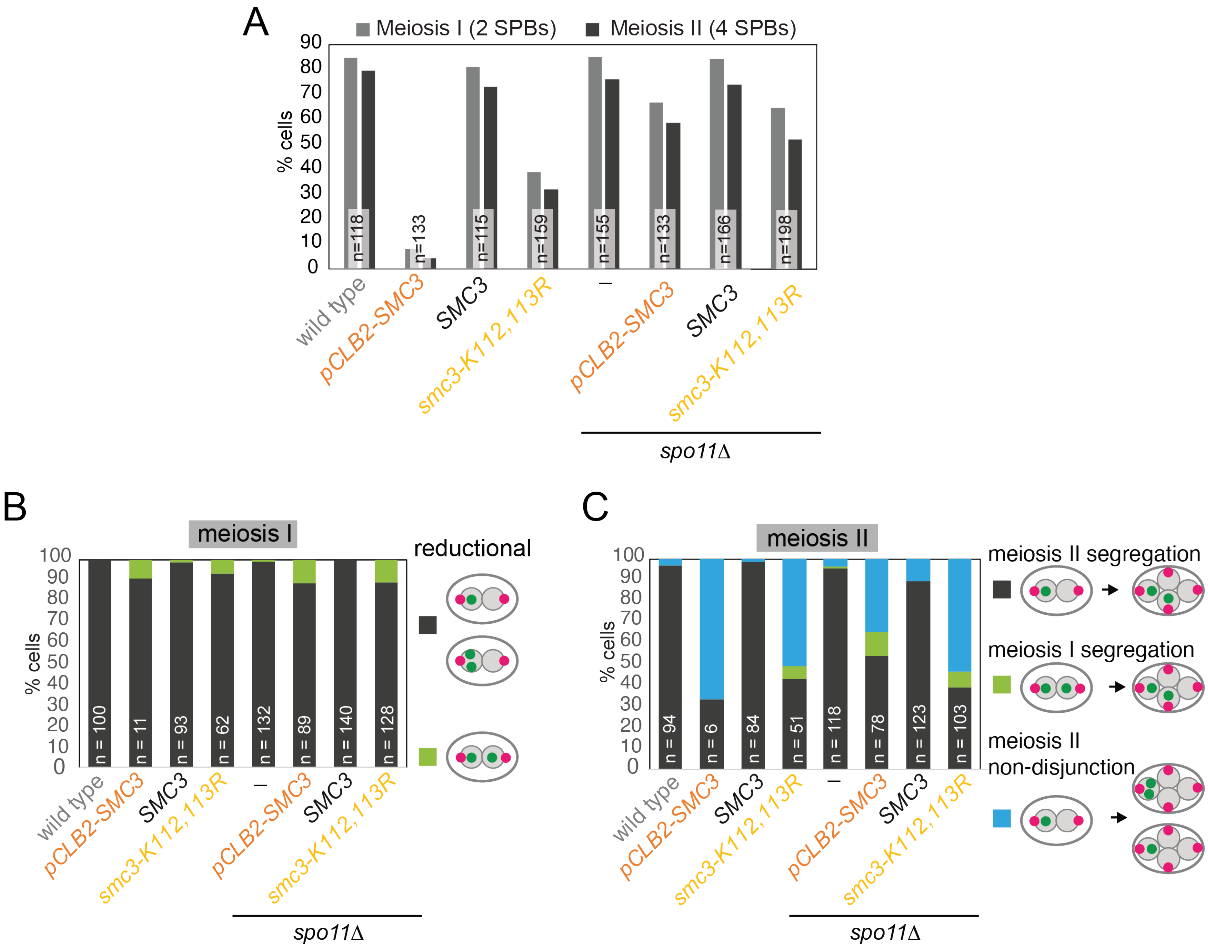


**Supplemental Figure S7. Prevention of meiotic recombination allows *smc3-K112,113R* cells to efficiently exit prophase.**

Live-cell imaging of *smc3-K112,113R* cells demonstrates that defective meiosis I sister chromatid monoorientation and meiosis II sister chromatid non-disjunction occur also in recombination-proficient cells that overcome the prophase delay, and that deletion of *SPO11* allows cells to overcome this delay. Cells of the indicated genotypes were grown and analyzed as described in Figure 6 (B-E). (A) Meiotic progression was scored based on the number of Spc42-tdTomato foci. (B) Meiosis I chromosome segregation (heterozygous *CENV*-GFP) was scored in cells with two separated Spc42-tdTomato foci. (C) Meiosis II chromosome segregation (heterozygous *CENV*-GFP) was scored in cells with four separated Spc42-tdTomato foci). Strains used were AM29317 (wild type), AM29318 (*pCLB2-3HA-SMC3*) AM29319 (*SMC3*), AM29320 (*smc3-K112,113R*), AM30238 (*spo11Δ*), AM30240 (*spo11Δ pCLB2-3HA-SMC3*), AM30242 (*spo11Δ SMC3*) and AM30244 (*spo11Δ smc3-K112,113R*).

### Supplemental Tables

**Table S1.** ***Saccharomyces cerevisiae*** **(SK1 or W303) and** ***Schizosaccharomyces pombe*** **strains used in this study**
Provided as an excel file.

**Table S2. List of plasmids used in this study**

| Plasmid | Description | Purpose and notes |
|---|---|---|
| AMp1342 | YIplac128-SMC3 | *LEU2* integration plasmid carrying *SMC3* |
| AMp1392 | YIplac128-smc3-K112R,K113R | *LEU2* integration plasmid carrying *smc3-K112R,K113R* |

### Table S3. List of qPCR primers used in this study

| Chr | Location | Distance from centromere | Primer Pair | Sequence | In figure |
|---|---|---|---|---|---|
| IV | Arm | -95kb | 782 | AGATGAAACTCAGGCTACCA | S5B-C |
| | | | 783 | TGCAACATCGTTAGTTCTTG | S5B-C |
| IV | Peri-centromere | -9.5kb | 1319 | ATGATTCAATGGATTTAGCC | S5B-C |
| | | | 1320 | GTCAGTCTTATGCTGTTCCC | S5B-C |
| IV | Centromere | +150bp | 794 | CCGAGGCTTTCATAGCTTA | S5B-C |
| | | | 795 | ACCGGAAGGAAGAATAAGAA | S5B-C |
| III | Centromere | -42bp | 8196 | ATAAACCAAACCCTTCCCCTTC | S5E |
| | | | 8197 | CCATATTGTTTGGCGCTGAT | S5E |
| IV | Arm | -95kb | 8175 | GCTACCACCAATAACACAGTTGAG | S5E |
| | | | 8176 | GTACCTTCCCTGATAATCCGTCT | S5E |
| IV | Centromere | +51bp | 8172 | GCCGAGGCTTTCATAGCTTA | S5E |
| | | | 8173 | GACGATAAAACCGGAAGGAAG | S5E |
| XII | Centromere | -45bp | 8206 | GGTTTGTAGACAACCAAACTGGTG | S5E |
| | | | 8207 | ACTCTTTACGCGGGTGTGTACT | S5E |

**Table S4. SK1 genome coordinates used to generate ChIP-Seq and Hi-C pile-up plots**

| Chr | Left arm peak | Left border | CEN | Right border | Right arm peak |
|---|---|---|---|---|---|
| I | 143,338 | 156,847 | 160,506 - 160,623 | 170,170 | 174,656 |
| II | 209,535 | 226,877 | 229,855 - 229,971 | 235,956 | 269,863 |
| III | 56,131 | 106,064 | 119,146 - 119,262 | 134,944 | 148,849 |
| IV | 412,039 | 452,984 | 461,938 - 462,053 | 468,978 | 191,167 |
| V | 143,196 | 150,521 | 156,659 - 156,776 | 170,642 | 194,898 |
| VI | 141,526 | 145,685 | 150,233 - 150,350 | 156,148 | 174,078 |
| VII | 480,232 | 500,759 | 508,233 - 508,352 | 515,930 | 563,982 |
| VIII | 72,321 | 95,832 | 100,974 - 101,091 | 105,373 | 114,788 |
| IX | 320,860 | 354,853 | 360,299 - 360,308 | 370,438 | 381,917 |
| X | 423,353 | 445,538 | 450,666 - 450,783 | 455,253 | 468,108 |
| XI | 417,329 | 438,536 | 445,725 - 445,841 | 452,317 | 477,072 |
| XIV | 132,607 | 137,781 | 154,749 - 154,867 | 165,549 | 185,746 |
| XIII | 249,427 | 254,063 | 266,751 - 266,870 | 278,483 | 293,988 |
| XIV | 592,647 | 625,567 | 637,885 - 638,002 | 656,081 | 660,807 |
| XV | 310,053 | 322,437 | 327,363 - 327,480 | 335,598 | 373,770 |
| XVI | 537,065 | 554,046 | 559,567 - 559,683 | 564,164 | 584,849 |

### Table S5. Hi-C libraries generated in this study

| Library name | Relevant genotype | Total R1/R2 aligned reads (M) | Valid unique Hi-C pairs (M) |
|---|---|---|---|
| HiC1_28719_wt | *RPL13A-2xFKBP12 fpr1Δ tor1-1 ndt80Δ* | 97,377,167/93,678,652 | 17,763,563 |
| *HiC1_28720_eco1-aa* | *RPL13A-2xFKBP12 fpr1Δ tor1-1 ndt80Δ ECO1-FRB-GFP* | 103,777,687/100,044,807 | 15,759,219 |
| HiC1_29750_wpl1 | *RPL13A-2xFKBP12 fpr1Δ tor1-1 ndt80Δ rad61Δ* | 77,203,756/74,233,462 | 13,525,545 |
| HiC1_29781_eco1-aa_wpl1 | *RPL13A-2xFKBP12 fpr1Δ tor1-1 ndt80Δ ECO1-FRB-GFP rad61Δ* | 95,353,060/92,175,659 | 11,292,250 |
| HiC2_11633_wt | *ndt80Δ* | 175,716,902/172,151,440 | 18,256,545 |
| HiC2_28841_clb5_clb6 | *ndt80Δ clb5Δ clb6Δ* | 160,012,048/156,826,193 | 20,128,179 |
